# Endogenous variation in ventromedial prefrontal cortex state dynamics during naturalistic viewing reflects affective experience

**DOI:** 10.1101/487892

**Authors:** Luke J. Chang, Eshin Jolly, Jin Hyun Cheong, Kristina M. Rapuano, Nathan Greenstein, Pin-Hao A. Chen, Jeremy R. Manning

**Affiliations:** Department of Psychological and Brain Sciences, Dartmouth College, Hanover, NH, 03755; Department of Psychology, Yale University, New Haven, CT 06511; Department of Psychology, National Taiwan University, Taipei, Taiwan

**Keywords:** vmPFC, endogenous processing, appraisal, affect

## Abstract

How we process ongoing experiences is shaped by our personal history, current needs, and future goals. Consequently, brain regions involved in generating these subjective appraisals, such as the vmPFC, often appear to be heterogeneous across individuals even in response to the same external information. To elucidate the role of the vmPFC in processing our ongoing experiences, we developed a computational framework and analysis pipeline to characterize the spatiotemporal dynamics of individual vmPFC responses as participants viewed a 45-minute television drama. Through a combination of functional magnetic resonance imaging, facial expression tracking, and self-reported emotional experiences across four studies, our data suggest that the vmPFC slowly transitions through a series of discretized states that broadly map onto affective experiences. Although these transitions typically occur at idiosyncratic times across people, participants exhibited a marked increase in state alignment during high affectively valenced events in the show. Our work suggests that the vmPFC ascribes affective meaning to our ongoing experiences.

## Introduction

Our brains process complex information with incredible speed and efficiency. This information can be broadly categorized into two distinct classes. First, our brains directly process *exogenous* information about the external environment by transducing physical phenomena (e.g., changes in energy, molecular concentrations, etc.) into sensory perceptions that allow us to generate and maintain a sense of what is happening around us (*1*, *2*). Mental representations that are directly driven by the external world are likely to be highly *similar* across individuals who share the same sensory experience. Second, our brains also process *endogenous* information that reflects our current internal homeostatic states, past experiences, and future goals (*3*). The integration of exogenous and endogenous information allows us to meaningfully interpret our surroundings, prioritize information that is relevant to our goals, and develop action plans (*4*). Given the same input information, individuals may have unique interpretations, feelings, and plans, often leading endogenous representations to be *idiosyncratic* across individuals. How can we establish a broad functional commonality across individuals when these specific endogenous experiences may be unique to each individual?

This conceptual distinction between exogenous and endogenous processes may reflect an organizing principle of the brain. Recent work characterizing the functional connectomes of human and macaque brains has revealed topographic gradients from cortical regions that process exogenous information originating from a single sensorimotor modality (e.g., auditory, visual, somatosensory/motor) to transmodal association cortex such as the default-mode network (DMN) (*5*, *6*). The DMN, which encompasses the ventromedial prefrontal cortex (vmPFC), posterior cingulate cortex (PCC), dorsomedial prefrontal cortex (dmPFC), and temporal parietal junction (TPJ) was initially identified to be metabolically active in the absence of any psychological task (*7*–*11*). Subsequently, the DMN has been linked to a number of endogenous processes such as: engaging in spontaneous internal thought and mind wandering (*12*–*14*), thinking about the self (*15*, *16*), prospecting into the future (*17*, *18*), and recalling autobiographical memories (*19*–*21*).

Within the DMN, vmPFC responses are particularly variable across individuals. Voxels in this region show little evidence of temporal synchronization across individuals when listening to auditory stories (*22*) or watching movies (*23*, *24*). Moreover, decoding accuracy using pattern classification is generally lower in the vmPFC compared to unimodal sensory areas (*25*), and functional connectivity patterns in resting state fMRI appear to be more variable across individuals in the vmPFC than in other areas of cortex (*26*, *27*).

Anatomically, the vmPFC projects directly to regions involved in affect and peripheral regulation, such as the hypothalamus, amygdala, ventral striatum, periaqueductal grey (PAG), and brainstem nuclei (*28*, *29*), as well as cognitive systems involved in conceptual processing such as the medial temporal lobe and dorsal and medial prefrontal cortex (*30*). With connections to systems involved in both affective and conceptual processing, the vmPFC is thought to be instrumental in making self-relevant evaluative appraisals (*3*, *4*, *15*), which are critical in generating feelings (*31*–*34*), evaluating the value of a choice option (*35*–*39*), and comprehending abstract narratives (*40*–*45*). The affective meaning generated by these appraisal processes are likely to be highly idiosyncratic across individuals.

Beyond an individual’s unique appraisals, there are several potential methodological factors contributing to the variability of vmPFC responses across individuals. First, the vmPFC is particularly vulnerable to susceptibility artifacts due to its close proximity to the orbital sinus, which can increase noise due to signal dropout and spatial distortions. These artifacts can be slightly mitigated by optimizing scanning parameters and sequences (*46*–*48*). Second, because the vmPFC is transmodal and integrates information from parallel processing streams, there may be greater variability in the temporal response profile of this region. While all brain regions exhibit some form of temporal integration, the length of temporal receptive windows are thought to be hierarchically organized based on levels of information processing (*49*, *50*). In early sensory regions, this integration occurs over short temporal receptive windows that reflect rapidly changing inputs. By contrast, regions involved in abstract processing (e.g., narrative comprehension) exhibit much longer temporal receptive windows (*51*–*53*). Together, this suggests that the heterogeneity observed in vmPFC signals arises from a combination of individual variability, spatial distortion, and temporal variation.

Establishing functional commonalities that transcend this variation in vmPFC response profiles requires identifying new ways to align signals across individuals. One such alignment procedure commonly used in the decision neuroscience literature employs an idiographic approach whereby experimental paradigms are customized to specific individuals (*37*, *54*–*56*). Based on an individual’s preferences, stimuli are mapped onto ordinal value signals, which are then used to identify brain regions that exhibit a linear correspondence (*57*). Another promising alignment approach attempts to remap regions across participants based on common temporal response profiles to the same time-locked external stimulation. Functional alignment algorithms such as hyperalignment (*58*, *59*) and the shared response model (*60*) can be highly effective at improving spatial alignment across people based on common functional responses while still maintaining individual differences (*61*). This approach might be particularly well suited for improving alignment in regions that process exogenous information and also potentially useful for mitigating variability arising from spatial distortions. However, because vmPFC responses reflect idiosyncratic appraisals based on endogenous information that are far removed from the exogenous input stimuli (e.g., goals, memories, etc), vmPFC activity patterns likely cannot be aligned across individuals in space or time using such approaches. Rather, aligning vmPFC responses across individuals might require developing a new approach that detects changes in latent states (e.g., mental, physiological).

In the present study, we investigate how the vmPFC processes endogenous information while viewing a rich naturalistic stimulus. Participants watched a 45-minute pilot episode of a character-driven television drama (*Friday Night Lights*) while undergoing functional magnetic resonance imaging (fMRI). This show was chosen in particular for its engaging story arc, parallel plotlines, and range of dynamics in affective experiences. We test the hypothesis that vmPFC activity reflects changes in latent states associated with the interaction between exogenous and endogenous processes. These states are hypothesized to reflect appraisals of meaningful plot elements as a narrative unfolds (e.g., “that is amazing!”, “that is terrible!”), that ultimately result in the emergence of feelings. More specifically, we predicted: (a) greater heterogeneity in spatiotemporal dynamics in vmPFC across individuals relative to primary sensory regions; (b) longer processing timescales in the vmPFC compared to primary sensory regions; and (c) that changes in vmPFC states correspond to meaningful psychological experiences.

## Results

### Idiosyncratic spatiotemporal dynamics of the vmPFC

We hypothesized that the vmPFC plays a critical role in processing endogenous information by integrating information from the external world with internal states, past experiences, and future goals. Unlike regions that directly process exogenous information (e.g. early visual cortex), we predicted that there should be little across-participant consistency in vmPFC responses during natural viewing. We tested this hypothesis using two separate studies. Participants in Study 1 (*n*=13) were scanned on a Philips 3T scanner while watching the first two episodes of *Friday Night Lights*. Participants in Study 2 (n=35) were scanned on a Siemens Prisma 3T scanner while watching only the first episode of the same show. First, we examined consistency in temporal responses across the 45-minute episode using inter-subject correlations (ISC; see *Methods*) (*23*). To reduce computational complexity, all analyses were performed on 50 non-overlapping regions based on a whole-brain parcellation using meta-analytic coactivations from over 10,000 published studies (*62*, *63*). We then computed the average pairwise temporal similarity of mean activity within each parcel across all participants. This yielded a number-of-subjects by number-of-subjects correlation matrix for each of the 50 parcels. ISC is simply the average of these pairwise similarities across participants. In Study 1, early sensory regions (e.g., V1) exhibited the highest level of temporal synchronization (mean ISC=0.38, sd=0.13, p < 0.001), whereas PFC parcels largely showed little evidence of synchronization across participants (mean vmPFC ISC=0.06, sd=0.07, p < 0.001; Figure 1A). These results were replicated in Study 2; V1 was associated with high levels of temporal ISC (mean ISC=0.35, sd=0.13, p < 0.001), whereas the vmPFC exhibited a low degree of synchronization (mean r=0.07, sd=0.05, p < 0.001; Figure 1B). The overall spatial pattern of ISC across brain parcels was highly similar across the two studies, r(48)=0.99, p < 0.001.

**Figure 1.**
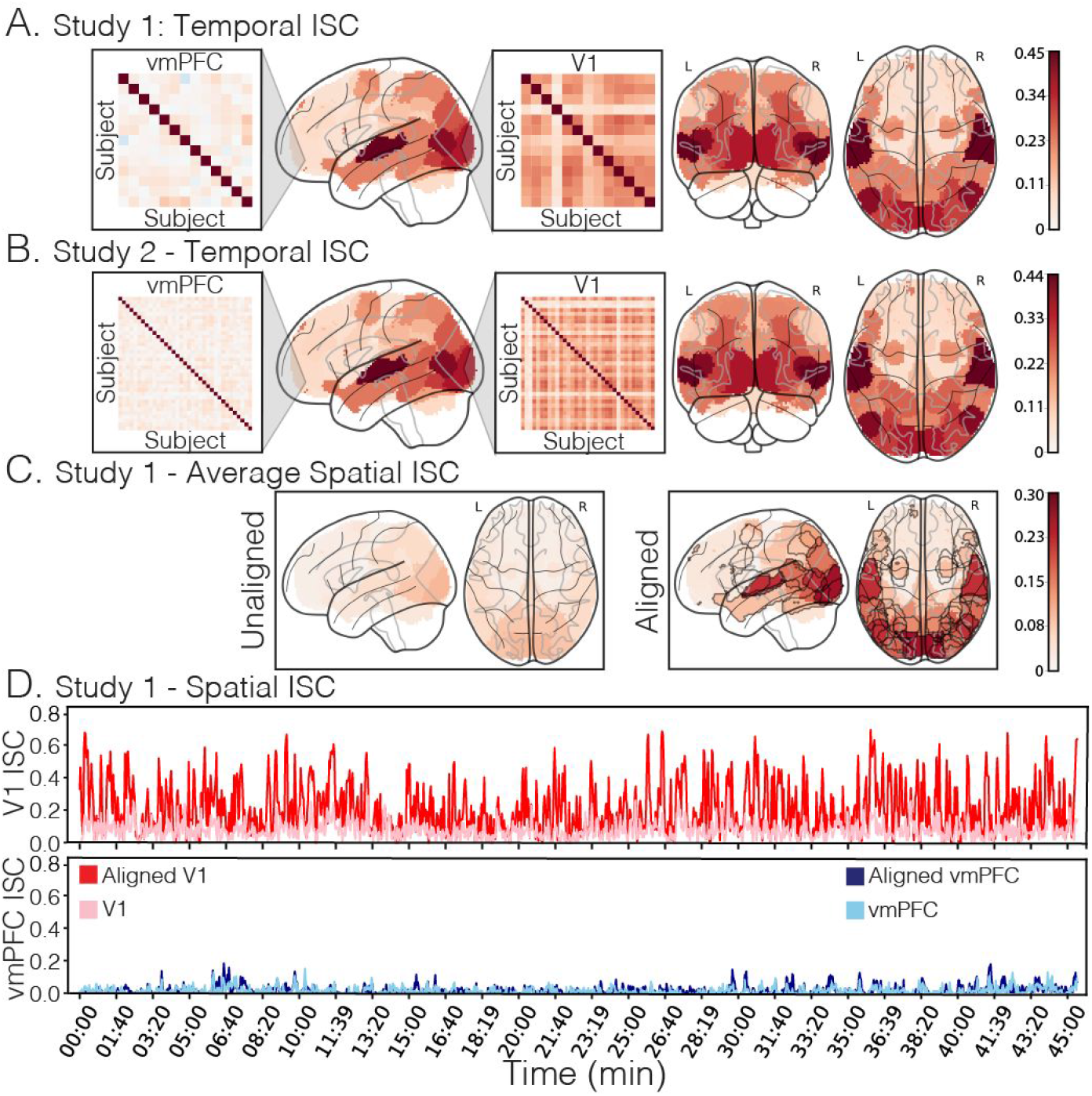
Intersubject correlations (ISC). A) Temporal ISC for each ROI for Study 1 (n=13). Temporal ISC reflects the average pairwise correlation between each individual participant’s activation time series while viewing the 45-minute episode. Heatmap blowouts depict all pairwise temporal ISCs between each participant for the vmPFC and V1 ROIs. B) Temporal ISC for each ROI for Study 2 (n=35); same format as Panel A. C) Spatial ISC across participants averaged across TRs for Study 1. Spatial ISC reflects the pairwise spatial similarity of activity patterns across participants within each ROI at each TR. Functional alignment in the right sub-panel was performed using the Shared Response Model (60) on an independent dataset. Parcels in which spatial ISC significantly increased following functional alignment using a permutation test are outlined in grey. D) Spatial ISC at each TR across the 45-minute episode for the vmPFC and V1 ROIs for Study 1. Lighter colors denote anatomical alignment of voxels across participants. Darker colors denote functional alignment.

Although vmPFC activity was idiosyncratic when considering the full duration of the episode, it is possible that activity might briefly align during particular moments of the episode. For example, an engaging pivotal moment in the story might evoke strong responses across individuals, whereas less salient moments might encourage person-specific endogenous processing. We defined a metric of instantaneous synchronization in Study 1 using the average spatial ISC computed across patterns of voxel activity within each ROI during each time point (i.e., each TR; see *Methods*). Unlike other methods of dynamic synchrony, this approach does not require specifying a sliding window length or shape (*64*) or restricting the analysis to a specific temporal frequency band (*65*). Instead, it characterizes the average alignment of spatial activity patterns across participants. Across all timepoints, we observed a modest spatial ISC in sensory regions, such as V1 (mean r=0.1, sd=0.06, Figure 1C) and virtually no synchronization in the vmPFC (mean r=0.01, sd=0.03, Figure 1C). One possible explanation for these findings is that spatial distortions from susceptibility artifacts might have increased variations in the spatial configuration of vmPFC voxels for each participant; or alternatively, that anatomical alignment techniques were insufficient for aligning fine-grain spatial patterns across individuals. For these reasons, we functionally aligned each region into a common 1000-dimensional space by fitting a Shared Response Model (*60*) to independent data (i.e., episode 2). Consistent with previous work, we found that functional alignment dramatically improved spatial synchronization across participants in regions involved in exogenous processing such as V1 (mean r=0.25, sd=0.19, p < 0.001, Figure 1D), but had no appreciable impact on vmPFC synchronization (mean r=0.02, sd=0.04, p=1.0, Figure 1D) (*58*–*60*).

These results demonstrate that vmPFC activity patterns do not appear to align spatially or temporally across people, which could arise from at least two possible explanations. First, it is possible that incoming stimulus-driven information does not directly impact vmPFC activity. For example, the vmPFC might play a role in representing or driving endogenous thoughts that are specifically *unrelated* to the exogenous experience (e.g., mind-wandering). Alternatively, the vmPFC may be involved in representing information or thoughts related to the stimulus (e.g., appraisals), but this information may be idiosyncratic to each individual.

### Temporal recurrence of spatial patterns

To more precisely characterize the structure of vmPFC responses across the episode, we examined the temporal recurrence of each individual’s’ unique patterns of vmPFC activity. For each participant, we computed a number-of-timepoints by number-of-timepoints correlation matrix reflecting the similarity of spatial patterns of vmPFC activity over the 45-minute episode. This analysis can aid in identifying time points during the show where a participant demonstrates a similar pattern of vmPFC activity, which might reflect a recurrence of a similar psychological state.

As shown in Figure 2A, this temporal correlation matrix exhibited a striking block diagonal structure. Specifically, the vmPFC exhibited periods of stable activity that persisted up to several minutes, punctuated by moments of rapid change (See Figs S1-S4 for all participants). This suggests that there are likely several different spatial configurations of vmPFC activity that are maintained for long periods of time. To determine if these response profiles are consistent across participants, we computed a one-sample *t*-test on Fisher *z*-transformed correlation values at each pair of time-points across all participants (Study 2, n=35). After correcting for multiple comparisons, we found very few time points that exhibited a consistent recurrence structure across participants (Figure 2B). Rather than treating each time point independently, we also examined whether the relative pattern of spatiotemporal dynamics was consistent across participants (i.e., the similarity of the lower triangles of the time x time spatial similarity matrix) (Figure 2C) and found little evidence of spatiotemporal synchronization (mean correlation=0.03, sd=0.01). In comparison, we observed a modest but reliable spatiotemporal synchronization across participants in V1 (mean correlation=0.12, sd=0.07, Figure 2C bottom) and a low spatial correlation in Posterior Cingulate Cortex (PCC; mean correlation=0.05, sd = 0.03; Figure S11B). Together, these results indicate that participants appear to share a block diagonal structure revealing long periods of sustained vmPFC activity, but the specific dynamics and patterns of recurrence appear to be idiosyncratic to each individual participant.

**Figure 2.**
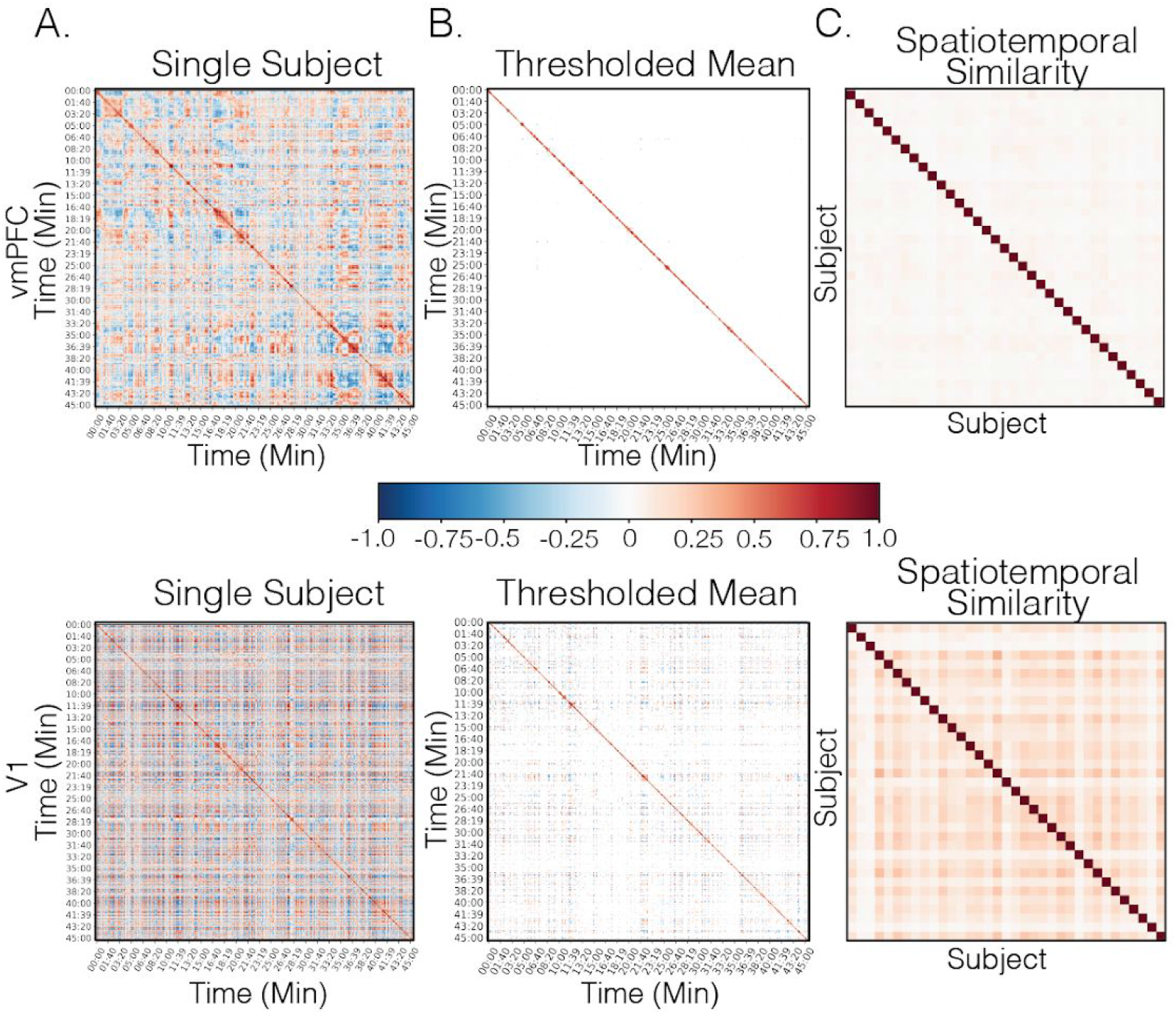
Temporal recurrence of spatial patterns. A) A recurrence matrix of vmPFC spatial patterns for a representative participant. B) A map of t-tests over each pair of time-points across all participants from Study 2 (n=35) thresholded using FDR q < 0.05 (vmPFC: p=0.0007; V1: p=0.011). C) A subject-by-subject similarity matrix representing the consistency of spatiotemporal recurrence patterns across participants. Colorbar is scaled between [-1,1], note that the scaling on the t-map depicted in panel B is one order of magnitude larger [-10,10].

### The vmPFC has a long temporal autocorrelation function

To quantify the rate of temporal integration of information in the vmPFC, we estimated the temporal autocorrelations of activity throughout the brain (*50*). Specifically, we estimated the amount of autocorrelation for each individual voxel in the brain up to 50 TR lags. We then fit an exponential function to the mean of these estimates within each ROI using a robust estimator and calculated the number of time-points before the function decayed to an arbitrary threshold of 0.1 (see *Methods*). Figure 3A illustrates the estimated amount of autocorrelation at the voxel level across each of the 50 parcels. Using a non-parametric sign test, we found that the autocorrelation in voxels in the vmPFC took longer to decay compared to V1 in both Study 1 (vmpfc: mean = 18.46 TR, sd=6.6 TR; v1: mean=3.62 TR, sd=1.19 TR, p < 0.001) and Study 2 (vmpfc: mean =8.46 TR, sd=2.95 TR; v1: mean=3.8 TR, sd=1.21 TR, p < 0.001). Additional supplementary analyses revealed that vmPFC voxels with longer autocorrelation were associated with a lower temporal signal to noise ratio (tSNR), indicating that this effect may be partially attributed to susceptibility artifacts (Study1: r=-0.72; Study2: r=-0.71). However, susceptibility artifacts could not fully account for our findings as we still observed a reliable distribution of voxel autocorrelation in the vmPFC across Study 1 and 2 after removing study-specific tSNR variance, r=0.58 (see Figure S5).

**Figure 3.**
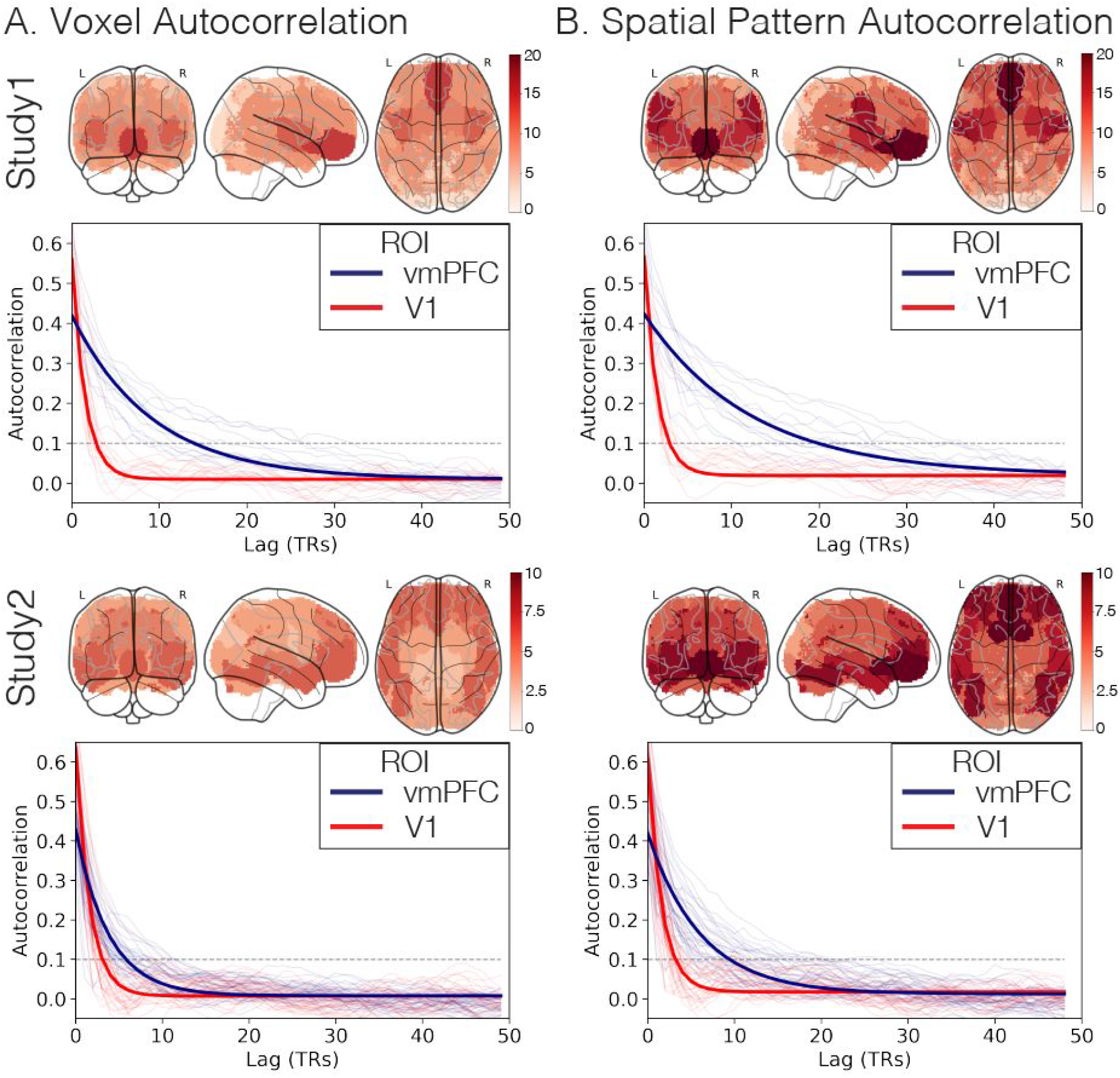
Temporal autocorrelation. A) The left panel displays the lag in TRs taken to reach an autocorrelation of 0.1 (arbitrarily selected). Larger values indicate more gradually changing voxel responses. In the right panel, we display the median autocorrelation function across voxels for vmPFC (blue) and V1 (red) for each participant. The darker lines reflect an exponential function fit to the median of the distribution across voxels for each ROI, and the thin lines represent autocorrelation functions for each individual participant. B) Identical analysis to Panel A using spatial patterns across all voxels within each ROI, rather than for individual voxels.

Next we sought to evaluate whether the multi-voxel spatial pattern within each ROI might be integrating information over a longer time window. We applied the same autocorrelation estimation procedure employed above using the multi-voxel pattern within each ROI. We found that the vmPFC displayed much more gradual changes in pattern responses compared to sensory regions such as V1 for both Study 1 (vmpfc: mean = 21.69 TR, sd=8 TR; v1: mean=4 TR, sd=1.47 TR, p < 0.001) and Study 2 (vmpfc: mean = 10.57 TR, sd=4.13 TR; v1: mean=4.06 TR, sd=1.63 TR, p < 0.001). Though we observed that the autocorrelation was longer for the vmPFC compared to V1 for both voxel and spatial patterns, only vmPFC patterns exhibited a significantly longer autocorrelation compared to the average voxel autocorrelation for both Study 1 (vmPFC: p=0.003; V1: p=0.12) and Study 2 (vmPFC: p<0.001; V1: p=0.08). This suggests that information encoded in spatial patterns in the vmPFC is being integrated over longer periods of time than sensory regions. Unlike previous work that has demonstrated similar findings by causally manipulating presented information at different temporal scales (e.g., scrambling words, sentences, or paragraphs) (*22*, *51*), this method allows us to estimate the longevity of spatial patterns within an ROI in a context-general way (including non stimulus-driven contexts) similar to exploring variations in power spectra (*52*, *53*, *66*, *67*).

### Spatial alignment of vmPFC latent states

The ISC analyses described above establish that the vmPFC does not show consistent responses across participants in its average time course or spatial patterns using either anatomical or functional alignment. Our temporal recurrence analysis revealed that each participant exhibited a similar block diagonal structure characterized by long periods of sustained vmPFC activity that recurred at multiple points throughout the episode. This suggests that the vmPFC may slowly transition between different states. Though the overall patterns of recurrence did not appear to synchronize across participants, it is possible that a subset of these states might be shared across participants, but expressed at different moments in time. To test this, we used Hidden Markov Models (HMMs) to segment patterns of activations in the vmPFC into discrete latent states. In implementing this analysis, we made several simplifying assumptions. We assumed that (a) voxel activations were drawn from orthogonal Gaussian distributions, (b) the transitions in latent states had the form of a first-order Markov chain, and (c) all participants experienced the same number of states. For each participant, we fit a separate HMM to a set of orthogonal principal components that explained 90% of the variance of the participants’ multivariate vmPFC time series. We used the Viterbi algorithm to determine the probability that the participant was in each given state at each moment of viewing the episode (Figure 4A) and aligned the states across participants by maximizing the spatial similarity using the Hungarian algorithm (*68*). To determine the number of states for the HMM estimation procedure, we identified the number of states that maximized the average within-state spatial similarity relative to the average between-state similarity. Overall, we found that a low number of states (k=4) appeared to yield the highest average within-state to between-state spatial similarity for vmPFC across both Study 1 and Study 2. We observed a slightly higher number of states for V1 for Study 1 (k=6) and Study 2 (k=7), and a large number of states for PCC for Study 1 (k=17), and for Study 2 (k=12) (Figure S9).

**Figure 4.**
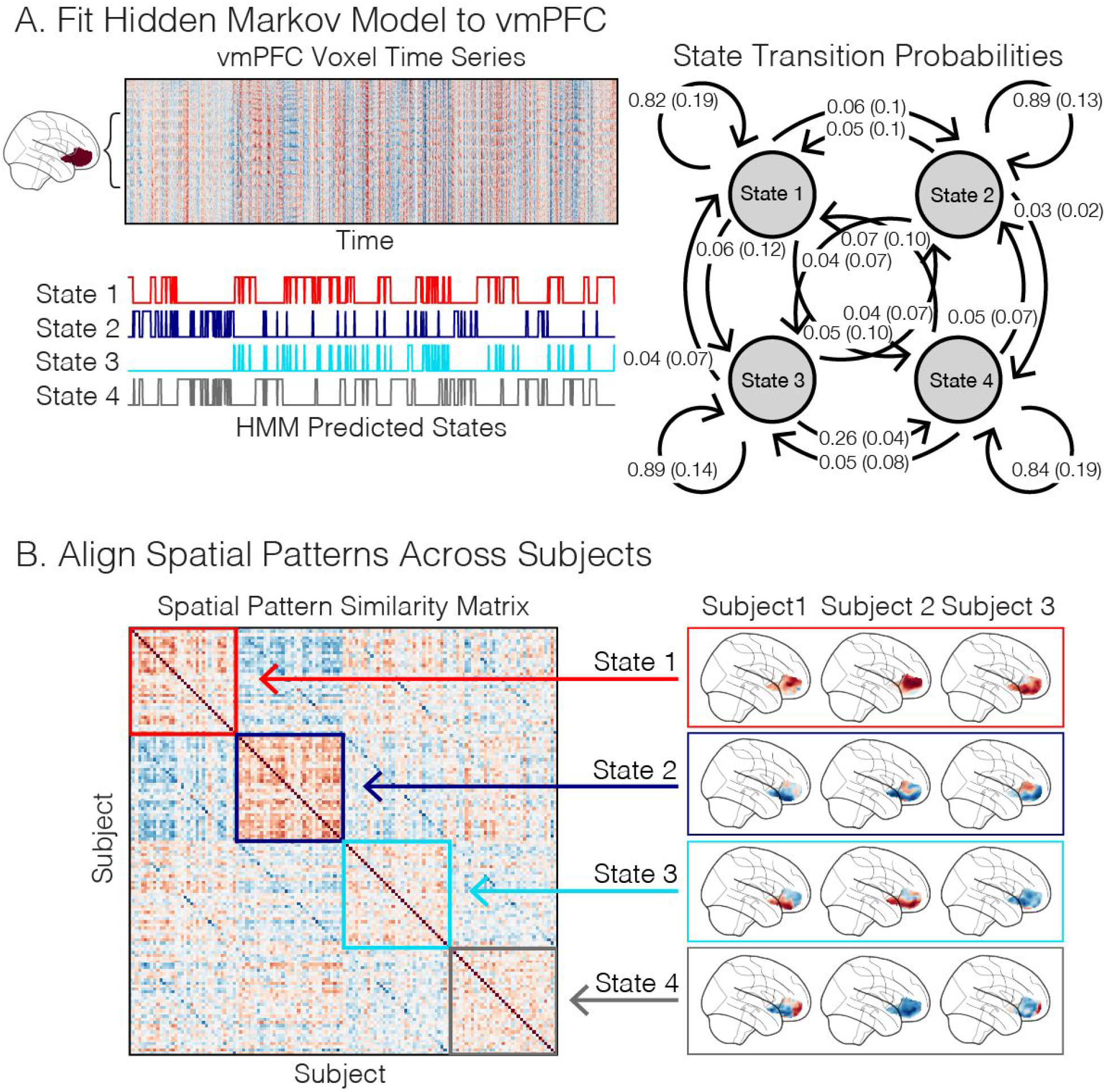
Individual-HMM estimation procedure. A) A schematic of how an HMM is fit to each participant’s ROI time series. We fit an HMM with a prespecified number of states to a PCA-reduced time series of voxel responses. This yielded estimates of the starting state, a transition matrix reflecting the likelihood of transitioning into any state at time t from time t-1, and a pattern of activity (assumed to be an emission from an orthogonal multivariate Gaussian distribution). We plot the average aligned transition probabilities for Study 2. We then applied the model to the same data to predict the most likely latent state at each moment in time using the Viterbi algorithm. B) We aligned states across participants by maximizing the spatial similarity of the gaussian emission weights associated with each estimated state using the Hungarian algorithm.

In contrast to the spatial ISC analyses, the Individual-HMM analysis revealed that a subset of the latent vmPFC states appeared to be shared across participants (Fig. 5A). The state with the highest spatial consistency in vmPFC and V1 are depicted in Figure 5B, and the corresponding spatial patterns averaged across participants are shown in Figure 5C. The magnitude of the HMM spatial similarity is in the upper range of what we observed in our spatial ISC analyses after performing functional alignment.

**Figure 5.**
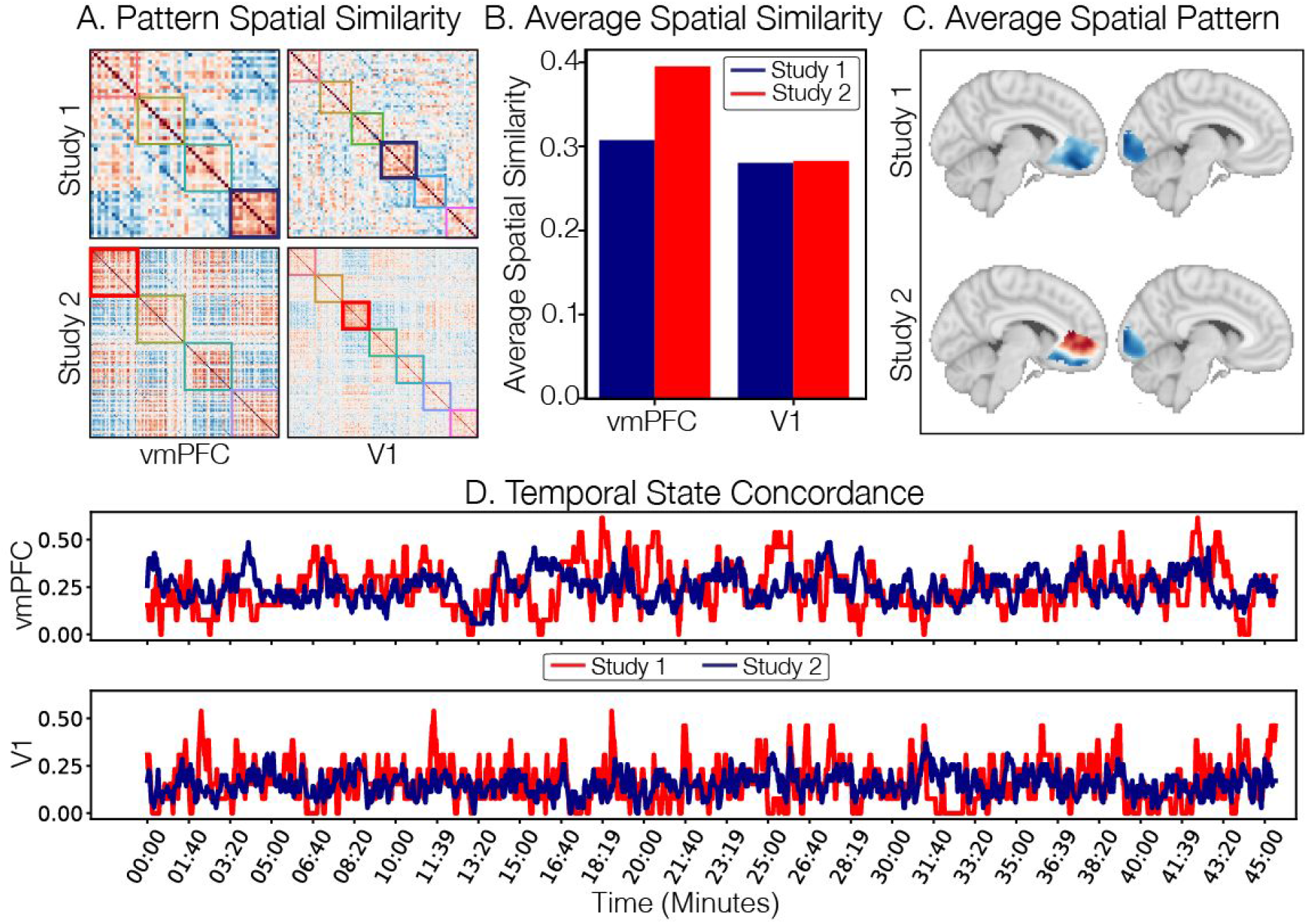
Alignment of spatial patterns corresponding to latent states. In this figure, we illustrate that the spatial patterns estimated from each individual HMM appear to be shared across participants. A) We plot the spatial similarity of the emission patterns corresponding to each individual’s state after aligning the states across participants using the Hungarian algorithm. The outlines indicate patterns associated with each participant for a given state in vmPFC, and V1 in Study 1 (n=13) and Study 2 (n=35). We highlight the vmPFC and V1 states with the highest intersubject spatial similarity (blue=Study 1, red=Study 2). B) We plot the magnitude of the spatial similarity for the states with the highest overall spatial similarity highlighted in panel A averaged across participants. C) We plot the vmPFC and V1 spatial patterns that correspond to the states associated with the highest intersubject spatial similarity from Study 1 and Study 2. These maps reflect the emission probabilities associated with each state averaged across participants. D) Temporal concordance of the highlighted predicted states across individuals within each study. Higher values indicate a higher proportion of participants occupying the same putative latent state within each study (blue: Study 1; red: Study 2).

Next we examined whether these latent states exhibited any temporal consistency using a concordance metric, which we defined as the proportion of participants occupying a given state at each moment of the episode. We plot the overall concordance of the vmPFC and V1 states that exhibited the highest spatial consistency across time in Figure 5D. This analysis indicates moderate state concordance in vmPFC (Study 1: mean=0.25, std=0.12; Study 2: mean=0.25, std=0.08), and V1 (Study 1: mean=0.16, std=0.11; Study 2: mean=0.15, std=0.06), but small levels of state concordance for PCC, presumably because of the larger number of states (Study 1 mean=0.05, std=0.06; Study 2 mean=0.08, std=0.04; Figure S11) (see Tables S1 and S2 for all states). The timecourses of these state concordances appear to show some degree of consistency across studies. We observed a moderate correlation in the temporal concordance between Studies 1 and 2 for V1, (r(1362)=0.24, p< 0.001), a small correlation for vmPFC (r(1362)=0.05, p = 0.07), and no significant correlation for PCC (r(1362)=-0.03, p = 0.23). Together, these analyses suggest that while it is possible to identify consistent spatial patterns in the vmPFC for specific states, these states tend not to be occupied by the same participants at the same moments in time (except at a few specific timepoints—i.e., local maxima in Figure 5D).

### Shared latent states using Group-HMM

The Individual-HMM analyses revealed that participants exhibited similar brain patterns for each latent state, but rarely occupied these states at the same moments in time. Building on these results, we estimated a Group-HMM for all participants in each study treating each participant as an independent sequence (*69*). This model simplifies our analysis pipeline and ensures that the voxel emission patterns associated with each hidden state are shared across participants within each study, but allows each participant to have a unique sequence of states. We selected the number of states in which the Bayesian Information Criterion (BIC; (*70*)) exhibited the largest improvement over a range of states (i.e., k=[2,25]; Figure S10). Similar to the analysis performed on each individual participant, this procedure identified a low number of states for both Study 1 and 2 (k=4). The transition probability matrices can be viewed in Figures S9.

We were additionally interested in the generalizability of the states across other contexts. We focused specifically on participants from Study 1 and trained an additional k=4 Group-HMM on the PCA-reduced vmPFC data from viewing episode 2. We aligned the states from the models trained on each episode using the Hungarian algorithm (*68*). We then assessed the cross-episode state concordance synchronization using Pearson correlations (Fig S8). Overall, we found that three of the four states appeared to be capturing similar states when testing the models on episode 1 (state1: r=0.3, p < 0.001; state 2: r=0.64, p < 0.001; state 3: r=1.0, p = 0.001; state 4: r=-0.03, p = 0.29; Figure S8A) and episode 2 (state1: r=0.3, p = 003; state 2: r=0.71, p = 0.002; state 3: r=1.0, p < 0.001; state 4: r=0.01, p = 0.93; Figure S8B) using a circle-shifting permutation test. These results provide evidence demonstrating that the vmPFC states estimated from a single viewing episode appear to generalize to other episodes.

### vmPFC state concordance is associated with affective experience

The above results demonstrate that participants exhibited similar spatial representations in the vmPFC based on the emission patterns associated with each latent HMM state. However, the time courses of how these patterns are expressed appear to be considerably less consistent across participants. We next sought to understand what types of thoughts or processes might be reflected by these vmPFC states.

We first performed a reverse correlation analysis (*23*) to identify which scenes from the show corresponded to the periods of high across-participant state concordance (Fig. 6A). We found that high concordance intervals occurred during scenes with strong narrative importance for the episode. For example, these key moments include scenes such as when the football players serve as mentors to young kids (~22 min), the protagonist is severely injured (~35 min), and the backup quarterback leads the team to a surprising victory despite the setbacks (~42 min). Further, these states appear to be maintained for durations on the order of multiple minutes, and are consistent across different participants and scanners.

**Figure 6.**
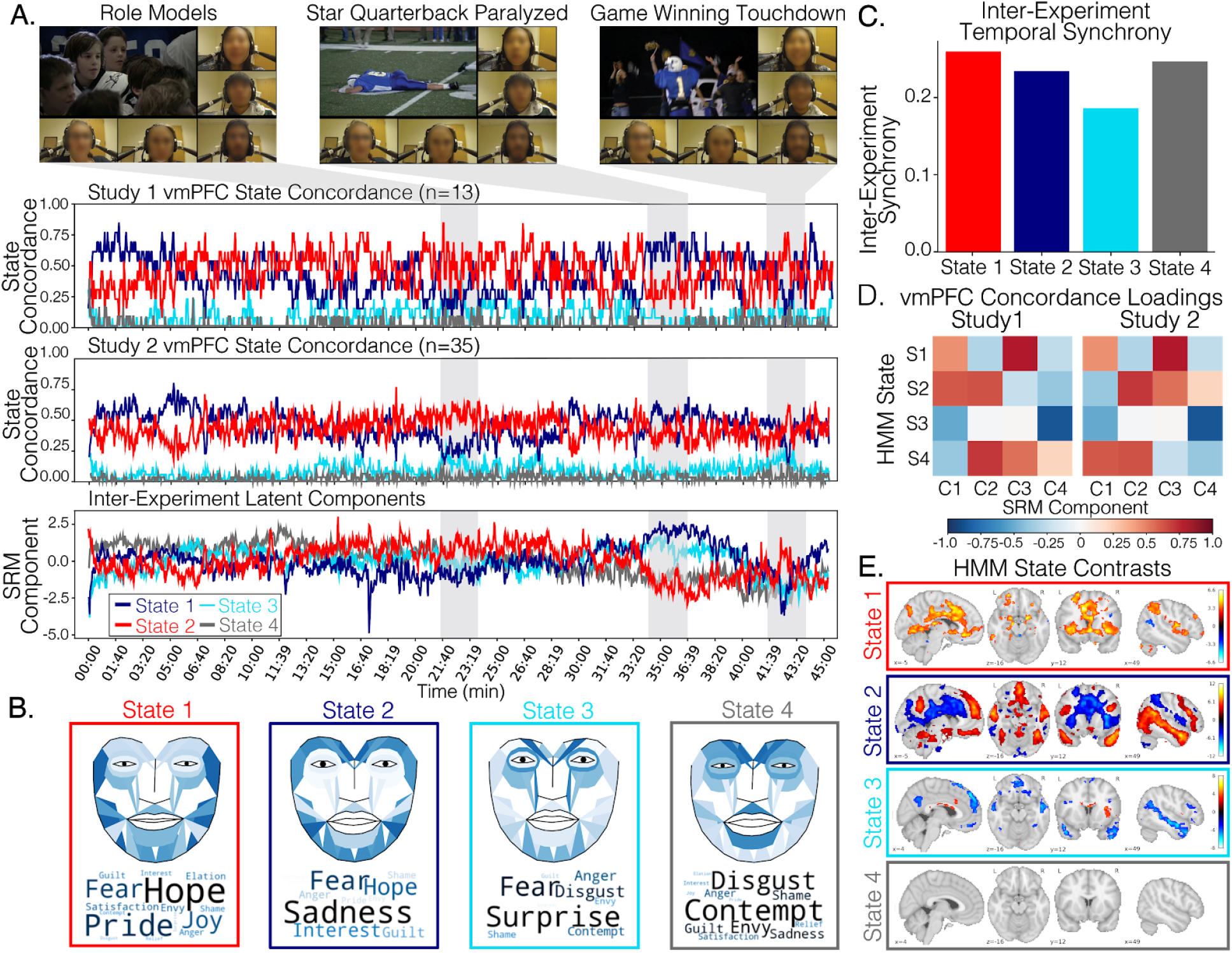
Affective experiences associated with increased state concordance. A) State concordance at each moment in time for each of the 4 states identified by the group vmPFC HMM (Studies 1 and 2). We also plot the inter-experiment latent components estimated using the Shared Response Model across these two studies, plus two additional behavioral studies (Study 3: Facial Expression) and subjective ratings (Study 4: Self-Reported Feelings). B) Here we visualize the transformation matrices that project Study 3 face expressions and Study 4 subjective feelings into the inter-experiment latent space. Darker colors in the face expressions indicate higher Action Unit intensity. Darker and larger words in the word cloud indicate higher contributions of the feeling onto the latent component. C) Overall inter-experiment temporal similarity for each estimated latent component across all 4 studies. D) Loadings of the vmPFC state concordances (Study 1 & Study 2) onto each inter-experiment latent component. E. Whole-Brain univariate contrasts based on vmPFC state occupancy. We computed the average voxel-wise activation associated with each state compared to all other states based and plot the group-level results thresholded with FDR q < 0.05.

To gain further quantitative insight into the identified vmPFC states, we collected two additional behavioral datasets to identify the facial expression behaviors and subjective self-reported feelings evoked by the television show in independent samples of participants. In Study 3, participants (n=20) watched the first episode of Friday Night Lights while their facial expressions were video recorded (*71*). We used a computer vision algorithm to identify 20 facial action units (AUs), a standardized system to describe the intensity of facial muscle movements (*72*), for each frame of the video during the episode (*73*). In Study 4, we recruited participants (n=188) from Amazon Mechanical Turk to watch the same television episode, which was periodically paused at 12 randomly selected timepoints spanning the episode, so that participants could report their subjective feelings on 16 different emotional dimensions. A different set of timepoints was chosen for each participant. We then used a collaborative filtering technique based on non-negative matrix factorization (see *Methods*) to predict how each participant was feeling for the remaining time points across all 16 dimensions.

We estimated a cross-experiment Shared Response Model (*74*) to identify a common latent space shared across all four experiments. This entailed learning study-specific transformation matrices that projected the following time series into a shared latent space: (1) the Study 1 four-dimensional vmPFC HMM state concordance time series, (2) the Study 2 four-dimensional vmPFC HMM state concordance time series, (3) the average Study 3 twenty-dimensional facial expression action unit time series, and (4) the average Study 4 sixteen-dimensional self-reported subjective emotion rating time series. The time course for each latent state was consistent across studies for all 4 states (mean ISC=0.23, std=0.03) and in the moderate range we observed for temporal ISC across the entire brain. The latent time series can be observed in Figure 6A and indicates a clear differentiation of signals at the three highlighted scenes. States 2 and 3 increase when the star quarterback experiences a traumatic spinal cord injury and undergoes emergency surgery (~35 min), while State 1 increases when the nervous and inexperienced backup quarterback throws an impressive pass that allows the team to come back and win the big game (~42 min). Importantly, these latent state time series capture the reliable changes in vmPFC HMM state concordance across both Study 1 and Study 2. However, we can also use the transformation matrices from Study 3 and Study 4 to directly interpret affective processes associated with these latent components (*75*). In Figure 6B, we project the transformation matrices back to action units to visualize the facial expressions displayed by participants in Study 3 at these particular time points (see also Figure S16). State 1 is most associated with increased intensity of AU12 (zygomaticus major; intensity=2.86), AU14 (Buccinator; intensity=3.36), and AU24 (Orbicularis oris; intensity=4), which presses the lips together and pulls the lip corners to form a smile (*76*). State 2, in contrast, is most associated with raising the eyebrows (AU1; Frontalis, pars medialis; intensity=3.11; & AU2; Frontalis pars lateralis; intensity=3.34), depressing the lip corners (AU15; Depressor anguli oris; intensity=3.09), raising the chin (AU17; Mentalis, intensity 3.27), and funneling the lips (AU18; Incisivii labii superioris; intensity=4.0). This suggests that vmPFC State 1 is associated with feelings of positive affect, whereas State 2 is associated with a feeling of surprise, concern, or worry. We visualized the transformation matrices from Study 4 using word clouds. Consistent with these interpretations of the facial expression behaviors, we found that the top five feelings associated with State 1 included Hope, Pride, Fear, Joy, and Satisfaction while the top five feelings associated with State 2 included Sadness, Fear, Hope, Interest, and Guilt (Figure S17).

Taken together, our results indicate that participants collectively experience at least two latent affective states while watching the episode, with our inter-experiment latent state model providing convergent evidence for distinct representations for positive and negative valence. When more participants from Studies 1 and 2 occupy State 1 based on their vmPFC patterns, participants in Studies 3 and 4 are smiling and experiencing positive feelings. In contrast, when more participants in Studies 1 and 2 occupy State 2 based on their vmPFC patterns, participants in Studies 3 and 4 are expressing and experiencing sadness and fear. This suggests that vmPFC states are linked to exogenous inputs and are more likely to align across individuals in contexts that elicit more intense affective responses (*4*, *33*, *77*).

These findings appear to be fairly specific to increased state synchronization in the vmPFC. vmPFC State 1 concordance increased at the same time points that participants smiled and subjectively reported feeling joy, while State 2 concordance increased when participants frowned and reported feeling sadness (Figure S13). V1 State concordance, in contrast, did not appear to be particularly coupled with changes in affect, but rather coupled strongly with large decreases in visual luminosity (Figure S12). Importantly, state concordance in other transmodal regions such as the PCC also did not appear to be strongly connected to affective experiences (Figure S14). More detailed feature mapping to time periods of high state concordance can be viewed in Figure S15.

### Whole-brain activations associated with each vmPFC state

To provide additional insight into the psychological states associated with each vmPFC state, we computed whole-brain univariate contrasts based on when the Group-HMM identified each participant’s vmPFC to be in a particular state. These analyses are identical to traditional task based univariate contrasts (*78*), but rather than identifying epochs based on manipulated experimental conditions, we used idiographic vmPFC state occupancy. In other words, we identified average brain activity that increased during time points when individual vmPFCs are in a specific state. Importantly, these time points vary across each individual, which means that these time points are locked to similar endogenous psychological states rather than exogenous events evoked by the stimulus. We employed meta-analytic decoding using the Neurosynth framework to provide quantitative reverse inferences for each map (*79*, *80*); see Figures S18). State 1 was associated with increased activity in the dorsal anterior cingulate (dACC), bilateral insula, bilateral, amygdala, and bilateral nucleus accumbens. These regions are associated with reward, fear, pain, motor, and somatosensory processing. State 2 was associated with increased activity in the vmPFC, PCC, dorsomedial prefrontal cortex (dmPFC), temporo-parietal junction (TPJ), and superior temporal sulcus (STS). These regions comprise the default mode and are associated with social processing. State 3 is associated with increased activity in the dACC and dorsal anterior insula. These regions are part of the salience network and are associated with executive control, attention, conflict, and learning. No voxels survived multiple comparisons correction for State 4. All contrasts are available on Neurovault (https://neurovault.org/collections/9062/).

## Discussion

In this study, we investigated how the vmPFC processes information while viewing a rich, evocative, multimodal naturalistic stimulus. We found evidence that the spatiotemporal response profiles of the vmPFC were heterogeneous across individuals, even after applying spatial and temporal alignment algorithms to the data. We also found that individual vmPFC spatial patterns appeared to persist for long periods of time and recurred periodically over the course of the episode. The long window of temporal integration in the vmPFC was present in individual voxels, but was enhanced in the spatial patterns. We used HMMs to segment patterns of vmPFC activity into discrete latent states and found that a subset of these states appeared to be shared across individuals. Although these states were most often expressed at different moments in time across individuals, scenes that evoked strong affective responses appeared to synchronize these latent states across participants, which replicated in an independent sample collected on a different scanner. Moreover, we observed converging evidence that these states synchronized with specific interpretable patterns of facial expressions and subjective affective ratings in two additional independent samples viewing the same television episode outside of the scanner. Taken together, these vmPFC activity states appear to reflect latent endogenous psychological processes involved in conferring affective meaning to ongoing events. This helps to explain the heterogeneity of responses within the vmPFC across people, and also has implications for experiments designed to probe the function of this region.

The vmPFC appears to be broadly involved in making meaning of the external world (*3*, *4*). It has been associated with representing and navigating cognitive maps and conceptual spaces (*81*–*83*) and interpreting narratives (*41*, *42*, *45*, *84*), which often requires abstracting temporal structure of abstract models of situations (*44*). At its most basic level, making a simple evaluation about the valence of an object or event (i.e., is it good or bad?) requires making an interpretation based on an individual’s past experiences, current homeostatic states, and future goals. This appraisal process appears to be directly linked to the vmPFC (*35*, *37*). For example, evaluating a particular food option requires integrating multiple features such as taste, cost, and caloric content (*85*) with broader goals such as eating healthy (*86*) and internal homeostatic drives (*87*). Beyond food, the vmPFC appears to code general representations of subjective valence appraisals that transcend stimulus modalities (*33*, *88*, *89*). Countless studies have implicated the vmPFC in processing the valence of emotional memories (*32*) and chemosensory sensory olfaction (*90*, *91*), and gustation (*56*) signals. These types of affective experiences of feelings, smells, and tastes can be causally induced with direct electrical stimulation of this region via intracranial electrodes implanted in patients with epilepsy (*34*). In this study, we find the highest level of synchronization of vmPFC states at the most emotionally evocative narrative events, which is consistent with work using pupillometry indicating that people temporally synchronize mental states at affectively arousing events (*77*).

Extracting meaning from our subjective experiences requires integrating multimodal information at a variety of timescales. Consistent with work that has demonstrated a cortical hierarchy of processing temporal information (*22*, *49*, *51*, *66*), we find that the vmPFC maintains stable states over timescales on the order of tens of seconds to minutes. While this slow processing is represented to some extent in the autocorrelations of individual voxels, it becomes more apparent when examining temporal autocorrelations of spatial patterns (*92*). Our autocorrelation approach complements experimental designs that shuffle information at multiple timescales (*22*, *51*). While most methods studying neural processes have focused on studying fast signals that are closer to the firing of individual neurons, fMRI may be particularly well suited for studying slower signals that involve integrating information across time such as the transformation of experience into memories (*24*, *93*).

Participants exhibited unique spatiotemporal response patterns in the vmPFC while viewing a common television episode. Our findings demonstrate that this cannot be solely accounted for by measurement issues (e.g., susceptibility artifacts, variations in hemodynamic response functions). This presents a significant challenge to traditional neuroimaging analysis methods that assume a common response profile across participants (e.g. two-level univariate analyses (*94*), resting state analyses (*95*), multivariate pattern analysis approaches (*96*, *97*), intersubject synchrony (*23*, *41*, *98*), and functional alignment (*58*–*60*, *98*)). Because subjective endogenous experiences are not typically shared across participants, regions like the vmPFC that appear to exhibit idiosyncratic stimulus-driven activity may be mischaracterized by these approaches. Our state-based analysis framework provides a means of characterizing this endogenous stimulus-driven activity, even when the response patterns do not align spatially or temporally across individuals, or to the external stimulus. Our approach may be also useful in translational applications where patient groups are often highly heterogeneous compared to healthy controls.

We note several limitations of our work. First, we assume that the anatomical demarcation of the vmPFC is consistent across all participants. We used a large ROI that was defined by a meta-analytic coactivation (*63*). However, the vmPFC can be further subdivided into more functionally specific regions (*29*), each which may have a unique response profile. We note that our pattern-based approach leverages variability across these subregions to identify state changes. Second, our HMM analyses assume that all participants experience the same number of latent states while viewing the television episode. This assumption was necessary in order to constrain the model and to evaluate spatial and temporal consistency across participants. However, it is unlikely that there is a ground truth “correct” number of states, or that all participants experience the same states. Finally, we do not directly link emotional experiences to vmPFC activity within a single participant. Instead, we provide converging evidence across different studies which are time-locked to the stimulus presentation. Video measurements of facial expressions while scanning is difficult due to occlusions from the head coil. However, simultaneous facial electromyography with fMRI has shown promise by demonstrating a relationship between displays of negative affect via facial expression and vmPFC activity (*99*) and may provide an avenue for future work.

In summary, we find evidence indicating heterogeneity in exogenous and endogenous information processing across the brain. The vmPFC appears to be involved in ascribing meaning to ongoing experiences with respect to the unique endogenous thoughts individuals bring to bear on processing each new moment. These processes may be reflected in spatial patterns (*79*, *100*–*103*), which appear to be unique to each appraisal and experience, and change at relatively slow timescales. We observed increased synchronization of these spatial patterns at emotionally salient scenes (*77*), suggesting that manipulating internal states (e.g., homeostatic states, memories, goals, or feelings) can increase vmPFC response synchrony across individuals. Our work demonstrates the potential of characterizing the temporal dynamics of spatial patterns to study the experiences of a single individual elicited during naturalistic multimodal scanning paradigms.

## Methods

### Study 1

#### Subjects

Thirteen subjects (*mean [sd] age = 24.61 [4.77]; 6 female*) were undergraduate and graduate students at Dartmouth College participating for either monetary compensation ($20/hr) or for partial course credit. All participants gave informed consent in accordance with the guidelines set by the Committee for the Protection of Human Subjects at Dartmouth College.

#### Procedure

Participants completed two separate sessions. In the first session, participants watched episodes 1 and 2 of the television drama *Friday Night Lights* (FNL) while undergoing functional Magnetic Resonance Imaging (fMRI). This show was chosen in particular for its engaging story arc, parallel plotlines, and range of affective experience dynamics. We scanned two separate episodes to ensure that we had sufficient data to perform functional alignment on independent data. Participants viewed each episode with a break in between followed by two runs of passively viewing images of characters. In a second session, participants watched episodes 3 and 4 outside of the scanner while we recorded their facial expressions using a head-mounted camera. All episodes were approximately 45-minutes in length. After each episode, participants made ratings of characters on a computer. The character viewing session, facial expressions of episodes 3 and 4, and the character ratings will be discussed in more detail in forthcoming manuscripts; those data are not reported in the present manuscript.

#### Imaging Acquisition

Data were acquired at the Dartmouth Brain Imaging Center (DBIC) on a 3T Philips Achieva Intera scanner (Philips Medical Systems, Bothell, WA) with a 32-channel phased-array SENSE (SENSitivity Encoding) head coil. Raw images were saved directly to NIfTI format. Structural images were acquired using a high-resolution T1-weighted 3D turbo field echo sequence: TR/TE: 8200/3.7ms, flip angle = 8°, resolution = 0.938 x 0.938 x 1.00 mm voxels, matrix size = 256 x 256, FOV = 240 x 240mm^2^. Functional blood-oxygenation-level-dependent (BOLD) images were acquired in an interleaved fashion using single-shot gradient-echo echo-planar imaging with pre-scan normalization, fat suppression and an in-plane acceleration factor of two (i.e. SENSE 2): TR/TE: 2000/30ms, flip angle = 75°, resolution = 3mm^3^ isotropic voxels, matrix size = 80 x 80, FOV = 240 x 240mm^2^, 40 axial slices with full brain coverage and no gap, anterior-posterior phase encoding. Functional scans were acquired during each episode in a single continuous run per episode (episode 1: 1364 TRs; episode 2: 1317 TRs).

### Study 2

#### Participants

Thirty-five participants (*mean [sd] age=19.0 [1.07] years; 26 female*) were recruited from undergraduate introductory psychology and neuroscience courses for either course credit or monetary compensation at Dartmouth College. All participants gave informed consent in accordance with the guidelines set by the Committee for the Protection of Human Subjects at Dartmouth College.

#### Procedure

The experimental procedure was identical to Study 1, with the exception that each participant only watched the first episode of FNL (45 minutes) while undergoing fMRI. This dataset provides a larger sample size on a different scanner to assess the replicability of our results.

#### Imaging Acquisition

Data were acquired at the Dartmouth Brain Imaging Center (DBIC) on a 3T Siemens Magnetom Prisma scanner (Siemens, Erlangen, Germany) with a 32-channel phased-array head coil. Raw DICOM images were converted to NIfTI images and stored in the brain imaging data structure (BIDS) format (*104*) using ReproIn from the ReproNim framework (*105*) ^95^. Structural images were acquired using a T1-weighted, single-shot, high-resolution MPRAGE sequence with an in-plane acceleration factor of two (i.e. GRAPPA 2) and pre-scan normalization: TR/TE: 2300/2.32ms, flip angle = 8°, resolution = 0.9mm^3^ isotropic voxels, matrix size = 256 x 256, FOV = 240 x 240mm^2^. Functional blood-oxygenation-level-dependent (BOLD) images were acquired in an interleaved fashion using gradient-echo echo-planar imaging with pre-scan normalization, fat suppression and an in-plane acceleration factor of two (i.e. GRAPPA 2), and no multiband (i.e. simultaneous multi-slice; SMS) acceleration: TR/TE: 2000/25ms, flip angle = 75°, resolution = 3mm^3^ isotropic voxels, matrix size = 80 x 80, FOV = 240 x 240mm^2^, 40 axial slices with full brain coverage and no gap, anterior-posterior phase encoding. All functional scans were acquired in a single continuous run (1364 TRs).

### Imaging Preprocessing

All MRI data for both studies 1 and 2 were preprocessed using a custom pipeline written using nipype (*106*). The pipeline involved trimming the first five non-steady-state TRs of functional each scan using nipy (*107*) and subsequently realigning each functional volume to the mean functional volume in a two-pass procedure implemented via FSL’s MCFLIRT (*108*). In parallel, brain-extraction (i.e. skull-stripping) was performed on T1-weighted structural images using ANTS (*109*). Subsequently, transformations for linearly coregistering realigned functional volumes to skull-stripped structural images, and non-linearly normalizing skull-stripped structural images to the ICBM 152 2mm MNI template were calculated using ANTS. These transforms were then concatenated and applied in a single step using basis spline interpolation in ANTS. Data were spatially smoothed using a 6mm FWHM Gaussian kernel implemented using fslmaths (*110*). To perform denoising, we fit a voxel-wise general linear model (GLM) for each participant using custom software implemented in Python (https://github.com/cosanlab/nltools). We used this toolbox to remove variance associated with the mean, linear, and quadratic trends, mean activity from a cerebral spinal fluid (CSF) mask, the effects of motion estimated during the realignment step using an expanded set of 24 motion parameters (6 demeaned realignment parameters, their squares, their derivatives, and their squared derivatives), motion spikes between successive TRs identified using ART (*111*) and global signal-intensity spikes greater than three standard-deviations above the mean intensity between successive TRs (Note: Study 1 did not include global signal intensity spikes). All analyses occured on the residual time-series after GLM estimation. Code for this custom pipeline is available at: https://github.com/cosanlab/cosanlab_preproc

#### Parcellation

All imaging analyses were conducted over a set of 50 parcels (parcellation available at http://neurovault.org/images/39711). The parcellation was created by performing a whole-brain parcellation of the coactivation patterns of activations across over 10,000 published studies available in the Neurosynth database (*62*, *63*). The use of a parcellation scheme has several advantages over the more conventional voxelwise and searchlight approaches. First, it is several orders of magnitude less computationally expensive as analyses are performed only 50 times compared to 352k voxels. Second, the parcels are non-overlapping and contain bilateral regions that reflect functional neuroanatomy, whereas a searchlight approach is limited to local spheres that do not adapt to different areas of cortex.

### Imaging Analyses

#### Intersubject Correlation

We take the intersubject correlation (ISC) to be the mean of the lower triangle of the pairwise correlation matrix of signals between each subject (see (*112*) for a tutorial).

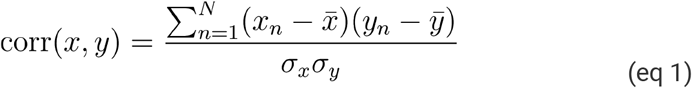

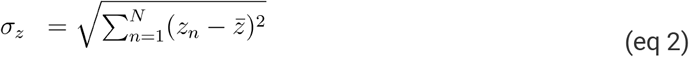

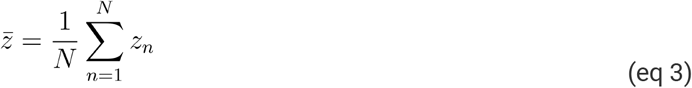

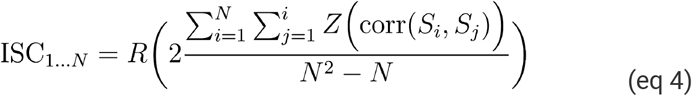

where

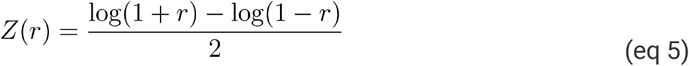

is the Fisher z-transformation and

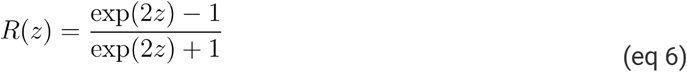

is its inverse.

*S* is a matrix of subject time series data from a total of *N* subjects and *i* and *j* indicate the subject indices. We performed hypothesis tests using a subject-wise bootstrap procedure as recommended by (*113*), which randomly draws participants with replacement to create a new sample by sample similarity matrix. Correlations between the same subject are dropped before computing ISC. We computed ISC for each subject’s mean vmPFC timeseries, their vmPFC spatial pattern at each moment in time, and their unique spatiotemporal pattern, defined as the vectorized lower triangle of the participant’s spatial recurrence matrix across all voxels in their vmPFC. We also carried out an analogous series of calculations with each participant’s V1 activity patterns. The similarity of spatiotemporal patterns has an interesting connection to computing distance correlation (*114*, *115*), a measure of general (i.e. possibly non-linear) statistical dependence between two signals of arbitrary dimensionality.

#### Functional Alignment

We performed functional alignment on Study 1 using the shared response model (SRM) (*60*). The deterministic SRM algorithm performs a joint PCA and can identify a reduced set of functional response components that maximally align across participants. We chose an arbitrarily selected 1000-dimensional embedding for each ROI. This algorithm learns a separate transformation matrix that projects each participant into this common space (*60*). Estimating a lower dimensional common model space can potentially aid in filtering out measurement noise that is assumed to be independent across individuals. To minimize bias, and ensure ISC analyses were estimated independent of computing the SRM (*58*); we learned transformation matrices using data from episode 2 and subsequently applied these transformations to data from episode 1. This allowed us to project each ROI’s voxel responses into the common 1000 dimensional space. To identify regions where spatial ISC significantly improved as a result of functional alignment, we performed a paired-sample sign permutation test on the dynamic ISC value using 5,000 samples and used a threshold of FDR q < 0.05 to correct for multiple comparisons.

#### Temporal Recurrence of Spatial Patterns

We created a spatial recurrence matrix by taking the pairwise Pearson correlation of the vectorized vmPFC activity for each TR collected during the 45-minute episode. This results in a number-of-TRs by number-of-TRs matrix (*n*=1,364 TRs) reflecting the degree to which the spatial activity patterns in the vmPFC are correlated at each pair of TRs. Sequential TRs where the spatial configuration of the vmPFC persists create a block diagonal structure. Recurring activity patterns at distant timepoints are reflected in the off-diagonal similarity.

#### Autocorrelation

To compute the autocorrelation function at each voxel, we calculated the average correlation of a voxel’s activation time series with itself shifted by a lag of *t* timesteps, for *t* ∈ {1, 2, …, 50}. This resulted in a correlation coefficient for each lag, for each voxel. We summarized the overall shape of this function within each ROI by taking the median across all voxels within the ROI and fitting a 3 parameter exponential function using a robust estimator. The parameter estimation was performed by minimizing the residual sum of squared error using the curve_fit() function from the scipy toolbox (*116*). We used the median of the distribution of autocorrelation across participants to ensure robust parameter estimation.

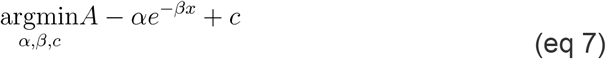

where *A* is the median autocorrelation function.

We used a similar procedure to calculate autocorrelation in spatial patterns. For each ROI, we computed a spatial recurrence matrix by correlating the activity patterns of each time *t* with the pattern at time *t* + *lag*, for *lag* ∈ {1, 2, …, 50}, and then averaging the matrices created for each timepoint. We fit an exponential function to the diagonal of the across-participants average spatial recurrence matrix.

To compare autocorrelations across ROIs, we calculated the number of TRs that the estimated exponential function took to reach an arbitrary correlation threshold of 0.1. This metric provides an intuitive and interpretable value for comparing the activity drift rates across regions and modalities (e.g. voxel versus spatial patterns) (*50*).

#### Hidden Markov Model

To identify state changes reflected in the neural dynamics within the vmPFC, we fit Hidden Markov Models (HMMs) to the voxel time series (*117*, *118*). HMMs are generative probabilistic models that propose that a sequence of observations is generated by a sequence of latent states. The transitions between these latent states are assumed to form a first-order Markov chain, such that the likelihood of transitioning to a new state from the current state can be predicted solely from the most recent data. We also assume that the observations are drawn from a multivariate Gaussian distribution with a diagonal covariance matrix. We performed two separate model fitting procedures: (1) an Individual-HMM where a model is fit separately to each participant’s data, (2) a Group-HMM where a single model is fit to all participants as a separate sequence.

First, we fit a separate HMM individually to each participant’s voxel pattern time series (vmPFC=3,591 voxels; V1=2,786 voxels). We reduced the dimensionality of the data using Principal Components Analysis (PCA) and retained enough components to explain 90% of the variance in the original signal (*66*). Reducing the dimensionality reduced computation time, decreased noise, and allowed us to better meet the orthogonality assumptions of the diagonal covariance matrix for the emission probabilities. The vmPFC had a higher number of components in both Study 1 (mean=58.34.43, sd=7.1) and Study 2 (mean=68.4.43, sd=15.74) compared to V1 in Study 1 (mean=33.69, sd=5.0) and Study 2 (mean=28.0, sd=6.42). We then fit the HMM using the Gaussian HMM function from the Python *hmmlearn* package (version 0.2.4; https://github.com/hmmlearn/hmmlearn), which estimates the model parameters using expectation-maximization. We used the fitted models to compute the viterbi sequence of states for each participant. We estimated a model for each subject over a range of k=[2,25]. For each model, we aligned the latent states of the participants to a randomly selected reference participant by maximizing the spatial correlation similarity between the estimated feature models using the Hungarian algorithm (*68*). We then selected the optimal value of k that maximized the average within state pattern spatial similarity relative to the between state pattern spatial similarity. This analysis demonstrated that subjects shared similar patterns of activity in vmPFC associated with each state, but these patterns were not expressed at the same moments in time.

Second, we fit a single Group-HMM model to all of the participants for each study by treating each participant’s voxel pattern time series as an independent sequence (*69*). This procedure learns a single voxel emission pattern for each state across all participants, but allows each participant to vary in the temporal sequence in the expression of the states. Because we cannot assess the spatial similarity of the emission patterns across participants as we did in the previous analysis, we selected the optimal number of states using the Bayesian Information Criterion (BIC; (*70*)). The BIC is a metric of model fit that penalizes the likelihood function for larger numbers of parameters to reduce overfitting defined as,

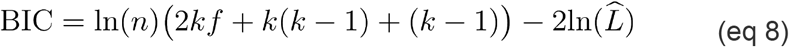

where *n* is the number of observations, *k* is the number of states, *f* is the number of features after reducing the dimensionality of the voxels using PCA, and 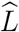 is the maximized log-likelihood of the HMM model given the data. Similar to the analyses above, we fit an HMM with Gaussian emissions assuming diagonal covariance to the principal components that explain 90% of the variance of the stacked participants’ vmPFC voxel time series (z-scored within each participant). This requires estimating 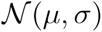 for each component, the transition probability matrix, and the initial starting probabilities. We selected the optimal states separately for each study by finding the largest decrease in BIC with respect to k, 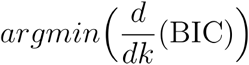.

#### Whole-Brain Univariate Group-HMM State Contrast

We used univariate contrasts to identify average activation in each voxel that was significantly different when participants occupied a particular state compared to all other states using the Group-HMM predictions. We used a one-sample t-test at the second-level to identify activity changes that are consistent across participants and used FDR q < 0.05 to identify a threshold that controls for multiple comparisons. This analysis is identical to traditional analyses used in task-based fMRI, except here we define different conditions based on individual state occupancy. Importantly, because each participant has a unique sequence of state transitions, this analysis identifies regions at the group level that are common across individuals but vary across time, which means that these regions are based on endogenous rather than exogenous processing.

#### HMM State Contrast Neurosynth Decoding

We used meta-analytic decoding (*79*) to make quantitative reverse inferences of the possible psychological functions implied by the pattern of brain activation using the Neurosynth framework (*119*). We used topic-based maps from previous studies (*80*, *120*, *121*), which were generated using Latent Dirichlet Allocation topic modeling of 9,204 fMRI articles and their corresponding coordinate data. Among the 80 topics, there were 48 related to psychological concepts, and we plot the top 10 that were most associated with each unthresholded Group-HMM State Contrast t-map based on spatial similarity using Pearson correlations in Figure S18.

### Study 3 (Face Expression)

#### Subjects

Twenty participants (mean [sd] age=18.9 [0.91] years; 13 female) were recruited from the Department of Psychological Brain sciences at Dartmouth College for course credit.

#### Procedure

Participants watched the first four episodes of the first season of Friday Night Lights over two separate two-hour sessions. Here we report results from episode one. Facial expressions during the experiment were monitored using GoPro HERO 4 cameras recording at 120 frames per second at 1920 x 1080 resolution. Each camera was positioned using a custom facecam headsets developed by our group (*71*). This approach is invariant to head motion and minimizes many types of facial occlusions. Recorded videos were then temporally aligned to the episodes by minimizing differences in audio intensity using our open source Python FaceSync toolbox version 0.0.8 (*71*). Facial behavioral features consisting of 20 facial action units (AU), a standard for measuring facial muscle movement based on the Facial Action Coding System (FACS, *72*) were extracted using the FACET algorithm (*73*) accessed through the iMotions biometric research platform (*122*). Data were downsampled to 0.5 hz.

#### Face Model Visualization

We used our Python Facial Expression Analysis Toolbox version 0.0.1 (*123*) to visualize how specific features from our model correspond to changes in facial morphometry data collected during Study 3. In brief, we learned a mapping between 20 facial action unit intensities and 68 landmarks comprising a 2-dimensional face using partial least squares implemented in scikit-learn (*124*). We used 10,708 images corresponding to each frame from the extended cohn-kanade facial expression database (CK+, *125*) and extracted landmarks using OpenFace (*126*) and action units from iMotions FACET engine (*122*). Next, we performed an affine transformation to a canonical face prior to fitting the model, and added pitch, roll, and yaw head rotation parameters as covariates with the 20 action unit features. We then fit our model using a 3-fold cross-validation procedure. We achieved an overall training model fit of r^2^ = .61 and a cross-validated mean of r^2^ = .53. We used this facial morphometry model to visualize the degree to which each AU loaded on a specific principal component. Action Unit intensities are overlaid onto corresponding facets of the face landmark model after performing a Min-Max feature scaling to ensure that the values are on the original AU scale [0,4], *x_scalsed_* = *std*(*x*)(4 – 0) + 0. Face expression loadings on the cross-experiment latent factor model can be viewed in Figure S16.

### Study 4 (Emotion Ratings)

#### Subjects

We recruited 192 participants from the Amazon Mechanical Turk workplace to participate in a 60-minute study (mean [sd] age=37.11 [10.71] years; 105 female), during which they viewed the first episode of Friday Night Lights. All participants provided informed consent in accordance with the guidelines set by the Committee for the Protection of Human Subjects at Dartmouth College and were compensated for their participation ($9). Four subjects were excluded for not having enough data to perform collaborative filtering (less than 25% of samples).

#### Procedure

The experimental paradigm was delivered via a custom open source web application built using Flask (http://flask.pocoo.org/), jsPsych (https://www.jspsych.org/), psiTurk (https://psiturk.org/), and MySQL (https://www.mysgl.com/) and served using an NGNIX (https://www.nginx.com/) web server hosted in our laboratory (https://github.com/cosanlab/moth_app, https://github.com/cosanlab/moth_turkframe). For each participant, the episode was paused over the course of its 45-minute runtime at random intervals of 200 - 280 seconds sampled from a uniform distribution (following an initial uninterrupted viewing period of 30 - 270 seconds). At each pause, participants were instructed to rate the intensity, if any, of 16 different dimensions: contempt, surprise, relief, anger, envy, shame, interest, elation, satisfaction, guilt, sadness, hope, pride, fear, joy, and disgust. The set of dimensions rated by each participant were presented in random order across participants. On average, participants completed 11.45 ratings (average sd=1.93 across emotion categories). The average inter-trial interval for making a rating was 229.65 sec (average sd = 53.33). Ratings were downsampled to 1 Hz and used to create a subject-by-time sparse matrix (188 x 2702). A collaborative filtering technique (non-negative matrix factorization with stochastic gradient descent) was applied to resulting matrices in order to infer each participant’s rating for every moment in time (*127*). Subjective emotion rating loadings on the cross-experiment latent factor model can be viewed in Figure S17.

#### Collaborative Filtering

We used collaborative filtering (CF) to create a timeseries for each affective dimension for every Study 4 participant (*128*). CF is a machine-learning technique that is commonly used by industrial recommender systems (e.g., Netflix, Amazon) to infer user preferences on many items (e.g., movies, music, products) when their preferences are only known for a few items (*129*). The key assumption driving CF is that, to the extent that participants *A* and *B* have similar ratings on *n* samples, they will rate other items similarly. To extend this general approach to the timeseries affective rating data we collected in Study 4, we convolved each participant’s (sparse) affective ratings (i.e., on a subset of the dimensions we sought to estimate) with a 60-second boxcar function kernel and divided each time point by the number of rating overlaps so that the interpolation reflects the mean rather than the sum of the ratings. This allows us to augment our data by assuming that time points closer in time are more similar to each other than time points further in time, but also acts as a low pass filter, which removes any fast high frequency changes that may be present in the data. Next, we used non-negative matrix factorization with stochastic gradient descent (100 iterations) implemented in our Python emotionCF toolbox (https://github.com/cosanlab/emotionCF) to perform the matrix completion (*127*). This approach assumes that there are N “types” of subjects that can be used as a basis set to linearly describe any individual subject, where N is the number of factors used in the matrix decomposition. The algorithm uses stochastic gradient descent to minimize the sum of squared error (with regularization) between the sparsely observed and predicted data. Because the dot product of the factorized matrices results in a full predicted matrix, this approach is able to make predictions for the unobserved data points for each participant. Importantly, this technique allows us to reconstruct a time series of each participant’s emotional ratings at every second during the show, while maintaining individual differences in rating time courses. Because we sparsely sample the subjective ratings, we believe this rating procedure should have a minimal impact on participants’ emotional experiences unlike alternative approaches that require dual-monitoring to continuously report feelings on single dimensions.

### Software

Unless otherwise noted, all of our imaging analyses were performed with our open source Python nltools toolbox version 0.4.2 (*130*) (https://github.com/cosanlab/nltools). This package wraps many scientific Python tools including: Nibabel 2.1 (*131*), nilearn 0.4 (*132*), numpy (*133*), pandas (*134*), scipy (*116*), and scikit-learn (*124*). Our plots were generated using matplotlib (*135*), seaborn (*136*), and custom plotting functions. All code used to perform the analyses in this paper are available on our lab github page (https://github.com/cosanlab/vmPFC_dynamics). Data will be shared on OpenNeuro upon publication of this manuscript.

## Acknowledgments

The authors would like to acknowledge Sushmita Sadhuka, Zainab Molani, and Antonia Hoidal who assisted in the data collection. This work was supported by funding from the National Institute of Mental Health R01MH116026 and R56MH080716 and NSF CAREER 1848370 as well as funding from the Young Scholar Fellowship Program by the Ministry of Science and Technology (MOST 109-2636-H-002-006) in Taiwan.

## Author Contributions

All authors were involved in conceptualizing this project. EJ & JC collected data for Study 1 & 2, JC collected data for Study 3, and NG designed and collected data for Study 4. EJ preprocessed Study 1 & 2, LC analyzed the data. All authors wrote the manuscript.

## Competing Interests

The authors declare no competing interests.

## Notes

### Competing Interest Statement

The authors have declared no competing interest.

### Summary of Updates

1) Added Group HMM Analyses. We have added a new Group-HMM that is fit to all participants data, treating each participant as a separate sequence. We believe this model complements our original analysis, and cleans up some of the noise associated with subject heterogeneity and simplifies our analysis as states are necessarily aligned across participants. We retain our original method to demonstrate the spatial similarity of the participant state patterns, but use the new Group-HMM for most of the other analyses. 2) Added cross-episode validation of Group-HMM. We have also fit the Group-HMM to Study 1 Episode 2 data and assess the generalizability of the patterns across episodes. Despite different narrative content, we see evidence that three of four states generalize across both episodes. 3) Added Inter-Experiment Latent Component Model. We have added an Inter-Experiment Shared Response Model, which allows us to identify patterns of facial expression and subjective feeling that project into a latent component that is shared with the vmPFC state concordances from Studies 1 & 2. We believe this analysis provides a more direct mapping of how latent affective components are manifested in vmPFC state concordances, face expressions, and subjective feelings. 4) Added additional feature mapping to HMM State Concordance. We provide additional analyses mapping the Group-HMM state concordances onto visual and affective features, which we believe helps to interpret what types of information the states may be processing. 5) Added additional ROI (PCC). We have added the Posterior Cingulate Cortex (PCC) as an additional control region for all analyses reported in the paper. 6) Deeper dive into autocorrelation. We have slightly updated our pipeline for computing autocorrelation and have included additional supplementary analyses exploring voxel-level autocorrelations. We now find that both voxels and spatial patterns in the vmPFC exhibit a longer autocorrelation compared to V1. In addition, we find that some of this variance can be explained by susceptibility artifact by mapping voxel autocorrelations onto signal-to-noise ratio maps. 7) Univariate contrasts based on vmPFC state changes. To provide additional evidence that the vmPFC states are processing distinct types of information, we have now included additional analyses in which we use participant-specific vmPFC state changes to identify differential voxel activations across the brain. These analyses indicate that three out of the four vmPFC states are associated with unique states reflected in whole-brain activity and highlight group level endogenous processing that is temporally offset from the stimulus.

## References

1. M. A. Goodale, A. D. Milner, Separate visual pathways for perception and action. Trends Neurosci. 15, 20–25 (1992).

2. M. Mishkin, L. G. Ungerleider, K. A. Macko, Object vision and spatial vision: two cortical pathways. Trends Neurosci. 6, 414–417 (1983).

3. Y. K. Ashar, L. J. Chang, T. D. Wager, Brain Mechanisms of the Placebo Effect: An Affective Appraisal Account. Annu. Rev. Clin. Psychol. 13, 73–98 (2017).

4. M. Roy, D. Shohamy, T. D. Wager, Ventromedial prefrontal-subcortical systems and the generation of affective meaning. Trends Cogn. Sci. 16, 147–156 (2012).

5. D. S. Margulies, S. S. Ghosh, A. Goulas, M. Falkiewicz, J. M. Huntenburg, G. Langs, G. Bezgin, S. B. Eickhoff, F. X. Castellanos, M. Petrides, E. Jefferies, J. Smallwood, Situating the default-mode network along a principal gradient of macroscale cortical organization. Proc. Natl. Acad. Sci. U. S. A. 113, 12574–12579 (2016).

6. J. M. Huntenburg, P.-L. Bazin, D. S. Margulies, Large-scale gradients in human cortical organization. Trends Cogn. Sci. 22, 21–31 (2018).

7. M. E. Raichle, A. M. MacLeod, A. Z. Snyder, W. J. Powers, D. A. Gusnard, G. L. Shulman, A default mode of brain function. Proc. Natl. Acad. Sci. U. S. A. 98, 676–682 (2001).

8. M. D. Greicius, B. Krasnow, A. L. Reiss, V. Menon, Functional connectivity in the resting brain: a network analysis of the default mode hypothesis. Proc. Natl. Acad. Sci. U. S. A. 100, 253–258 (2003).

9. G. L. Shulman, J. A. Fiez, M. Corbetta, R. L. Buckner, F. M. Miezin, M. E. Raichle, S. E. Petersen, Common blood flow changes across visual tasks: II. decreases in cerebral cortex. J. Cogn. Neurosci. 9, 648–663 (1997).

10. D. A. Gusnard, E. Akbudak, G. L. Shulman, M. E. Raichle, Medial prefrontal cortex and self-referential mental activity: relation to a default mode of brain function. Proc. Natl. Acad. Sci. U. S. A. 98, 4259–4264 (2001).

11. M. D. Fox, A. Z. Snyder, J. L. Vincent, M. Corbetta, D. C. Van Essen, M. E. Raichle, The human brain is intrinsically organized into dynamic, anticorrelated functional networks. Proc. Natl. Acad. Sci. U. S. A. 102, 9673–9678 (2005).

12. J. R. Andrews-Hanna, J. Smallwood, R. N. Spreng, The default network and self-generated thought: component processes, dynamic control, and clinical relevance. Ann. N. Y. Acad. Sci. 1316, 29–52 (2014).

13. M. F. Mason, M. I. Norton, J. D. Van Horn, D. M. Wegner, S. T. Grafton, C. N. Macrae, Wandering minds: the default network and stimulus-independent thought. Science. 315, 393–395 (2007).

14. K. Christoff, A. M. Gordon, J. Smallwood, R. Smith, J. W. Schooler, Experience sampling during fMRI reveals default network and executive system contributions to mind wandering. Proc. Natl. Acad. Sci. U. S. A. 106, 8719–8724 (2009).

15. W. M. Kelley, C. N. Macrae, C. L. Wyland, S. Caglar, S. Inati, T. F. Heatherton, Finding the self? An event-related fMRI study. J. Cogn. Neurosci. 14, 785–794 (2002).

16. A. C. Jenkins, C. N. Macrae, J. P. Mitchell, Repetition suppression of ventromedial prefrontal activity during judgments of self and others. Proc. Natl. Acad. Sci. U. S. A. 105, 4507–4512 (2008).

17. D. L. Schacter, D. R. Addis, R. L. Buckner, Remembering the past to imagine the future: the prospective brain. Nat. Rev. Neurosci. 8, 657–661 (2007).

18. R. L. Buckner, D. C. Carroll, Self-projection and the brain. Trends Cogn. Sci. 11, 49–57 (2007).

19. R. Cabeza, P. St Jacques, Functional neuroimaging of autobiographical memory. Trends Cogn. Sci. 11, 219–227 (2007).

20. R. N. Spreng, R. A. Mar, A. S. N. Kim, The common neural basis of autobiographical memory, prospection, navigation, theory of mind, and the default mode: a quantitative meta-analysis. J. Cogn. Neurosci. 21, 489–510 (2009).

21. J. Rissman, H. T. Greely, A. D. Wagner, Detecting individual memories through the neural decoding of memory states and past experience. Proc. Natl. Acad. Sci. U. S. A. 107, 9849–9854 (2010).

22. Y. Lerner, C. J. Honey, L. J. Silbert, U. Hasson, Topographic mapping of a hierarchy of temporal receptive windows using a narrated story. J. Neurosci. 31, 2906–2915 (2011).

23. U. Hasson, Y. Nir, I. Levy, G. Fuhrmann, R. Malach, Intersubject synchronization of cortical activity during natural vision. Science. 303, 1634–1640 (2004).

24. J. Chen, Y. C. Leong, C. J. Honey, C. H. Yong, K. A. Norman, U. Hasson, Shared memories reveal shared structure in neural activity across individuals. Nat. Neurosci. 20, 115–125 (2017).

25. A. Bhandari, C. Gagne, D. Badre, Just above Chance: Is It Harder to Decode Information from Human Prefrontal Cortex Blood Oxygenation Level-dependent Signals? J. Cogn. Neurosci., 1–26 (2018).

26. S. Mueller, D. Wang, M. D. Fox, B. Yeo, J. Sepulcre, Individual variability in functional connectivity architecture of the human brain. Neuron (2013).

27. E. M. Gordon, T. O. Laumann, B. Adeyemo, J. F. Huckins, W. M. Kelley, S. E. Petersen, Generation and evaluation of a cortical area parcellation from resting-state correlations. Cereb. Cortex. 26, 288–303 (2016).

28. S. N. Haber, K. Kunishio, M. Mizobuchi, E. Lynd-Balta, The orbital and medial prefrontal circuit through the primate basal ganglia. J. Neurosci. 15, 4851–4867 (1995).

29. T. Kahnt, L. J. Chang, S. Q. Park, J. Heinzle, J.-D. Haynes, Connectivity-based parcellation of the human orbitofrontal cortex. J. Neurosci. 32, 6240–6250 (2012).

30. J. R. Andrews-Hanna, J. S. Reidler, J. Sepulcre, R. Poulin, R. L. Buckner, Functional-anatomic fractionation of the brain’s default network. Neuron. 65, 550–562 (2010).

31. A. R. Damasio, Descartes’ error (Random House, 2006).

32. A. R. Damasio, T. J. Grabowski, A. Bechara, H. Damasio, L. L. Ponto, J. Parvizi, R. D. Hichwa, Subcortical and cortical brain activity during the feeling of self-generated emotions. Nat. Neurosci. 3, 1049–1056 (2000).

33. J. Chikazoe, D. H. Lee, N. Kriegeskorte, A. K. Anderson, Population coding of affect across stimuli, modalities and individuals. Nat. Neurosci. 17, 1114–1122 (2014).

34. K. C. R. Fox, J. Yih, O. Raccah, S. L. Pendekanti, L. E. Limbach, D. D. Maydan, J. Parvizi, Changes in subjective experience elicited by direct stimulation of the human orbitofrontal cortex. Neurology. 91, 1–9 (2018).

35. A. Rangel, C. Camerer, P. R. Montague, A framework for studying the neurobiology of value-based decision making. Nat. Rev. Neurosci. 9, 545–556 (2008).

36. C. Padoa-Schioppa, J. A. Assad, Neurons in the orbitofrontal cortex encode economic value. Nature. 441, 223–226 (2006).

37. H. Plassmann, J. O’Doherty, B. Shiv, A. Rangel, Marketing actions can modulate neural representations of experienced pleasantness. Proc. Natl. Acad. Sci. U. S. A. 105, 1050–1054 (2008).

38. T. Kahnt, J. Heinzle, S. Q. Park, J.-D. Haynes, The neural code of reward anticipation in human orbitofrontal cortex. Proc. Natl. Acad. Sci. U. S. A. 107, 6010–6015 (2010).

39. J. P. O’Doherty, M. L. Kringelbach, E. T. Rolls, J. Hornak, C. Andrews, Abstract reward and punishment representations in the human orbitofrontal cortex. Nat. Neurosci. 4, 95–102 (2001).

40. E. C. Ferstl, J. Neumann, C. Bogler, D. Y. Von Cramon, The extended language network: a meta-analysis of neuroimaging studies on text comprehension. Hum. Brain Mapp. 29, 581–593 (2008).

41. E. Simony, C. J. Honey, J. Chen, O. Lositsky, Y. Yeshurun, A. Wiesel, U. Hasson, Dynamic reconfiguration of the default mode network during narrative comprehension. Nat. Commun. 7, 12141 (2016).

42. L. J. Silbert, C. J. Honey, E. Simony, D. Poeppel, U. Hasson, Coupled neural systems underlie the production and comprehension of naturalistic narrative speech. Proc. Natl. Acad. Sci. U. S. A. 111, E4687–96 (2014).

43. A. G. Huth, W. A. de Heer, T. L. Griffiths, F. E. Theunissen, J. L. Gallant, Natural speech reveals the semantic maps that tile human cerebral cortex. Nature. 532, 453–458 (2016).

44. C. Baldassano, U. Hasson, K. A. Norman, Representation of real-world event schemas during narrative perception. J. Neurosci. 38, 9689–9699 (2018).

45. Y. Yeshurun, S. Swanson, E. Simony, J. Chen, C. Lazaridi, C. J. Honey, U. Hasson, Same Story, Different Story. Psychol. Sci. 28, 307–319 (2017).

46. N. Weiskopf, C. Hutton, O. Josephs, R. Turner, R. Deichmann, Optimized EPI for fMRI studies of the orbitofrontal cortex: compensation of susceptibility-induced gradients in the readout direction. Magnetic Resonance Materials in Physics, Biology and Medicine. 20, 39–49 (2007).

47. Y. P. Du, M. Dalwani, K. Wylie, E. Claus, J. R. Tregellas, Reducing susceptibility artifacts in fMRI using volume-selective z-shim compensation. Magn. Reson. Med. 57, 396–404 (2007).

48. G. H. Glover, C. S. Law, Spiral-in/out BOLD fMRI for increased SNR and reduced susceptibility artifacts. Magn. Reson. Med. 46, 515–522 (2001).

49. U. Hasson, J. Chen, C. J. Honey, Hierarchical process memory: memory as an integral component of information processing. Trends Cogn. Sci. 19, 304–313 (2015).

50. J. D. Murray, A. Bernacchia, D. J. Freedman, R. Romo, J. D. Wallis, X. Cai, C. Padoa-Schioppa, T. Pasternak, H. Seo, D. Lee, X.-J. Wang, A hierarchy of intrinsic timescales across primate cortex. Nat. Neurosci. 17, 1661–1663 (2014).

51. C. J. Honey, T. Thesen, T. H. Donner, L. J. Silbert, C. E. Carlson, O. Devinsky, W. K. Doyle, N. Rubin, D. J. Heeger, U. Hasson, Slow cortical dynamics and the accumulation of information over long timescales. Neuron. 76, 423–434 (2012).

52. A. T. Baria, M. N. Baliki, T. Parrish, A. V. Apkarian, Anatomical and functional assemblies of brain BOLD oscillations. J. Neurosci. 31, 7910–7919 (2011).

53. J.-P. Kauppi, I. P. Jääskeläinen, M. Sams, J. Tohka, Inter-subject correlation of brain hemodynamic responses during watching a movie: localization in space and frequency. Front. Neuroinform. 4, 5 (2010).

54. J. P. O’Doherty, P. Dayan, K. Friston, H. Critchley, R. J. Dolan, Temporal difference models and reward-related learning in the human brain. Neuron. 38, 329–337 (2003).

55. J. W. Kable, P. W. Glimcher, The neural correlates of subjective value during intertemporal choice. Nat. Neurosci. 10, 1625–1633 (2007).

56. S. M. McClure, J. Li, D. Tomlin, K. S. Cypert, L. M. Montague, P. R. Montague, Neural correlates of behavioral preference for culturally familiar drinks. Neuron. 44, 379–387 (2004).

57. J. P. O’Doherty, A. Hampton, H. Kim, Model-based fMRI and Its application to reward learning and decision making. Ann. N. Y. Acad. Sci. 1104, 35–53 (2007).

58. J. V. Haxby, J. S. Guntupalli, A. C. Connolly, Y. O. Halchenko, B. R. Conroy, M. I. Gobbini, M. Hanke, P. J. Ramadge, A common, high-dimensional model of the representational space in human ventral temporal cortex. Neuron. 72, 404–416 (2011).

59. J. S. Guntupalli, M. Hanke, Y. O. Halchenko, A. C. Connolly, P. J. Ramadge, J. V. Haxby, A model of representational spaces in human cortex. Cereb. Cortex. 26, 2919–2934 (2016).

60. P.-H. Chen, J. Chen, Y. Yeshurun, U. Hasson, J. Haxby, P. J. Ramadge, in Advances in Neural Information Processing Systems 28, C. Cortes, N. D. Lawrence, D. D. Lee, M. Sugiyama, R. Garnett, Eds. (Curran Associates, Inc., 2015), pp. 460–468.

61. F. Ma, S. A. Nastase, J. S. Guntupalli, J. V. Haxby, Reliable individual differences in fine-grained cortical functional architecture. Neuroimage. 183, 375–386 (2018).

62. T. Yarkoni, A. de la Vega, L. J. Chang, Fully automated meta-analytic clustering and decoding of human brain activity.

63. A. de la Vega, L. J. Chang, M. T. Banich, T. D. Wager, T. Yarkoni, Large-scale meta-analysis of human medial frontal cortex reveals tripartite functional organization. J. Neurosci. 36, 6553–6562 (2016).

64. A. S. Choe, M. B. Nebel, A. D. Barber, J. R. Cohen, Y. Xu, J. J. Pekar, B. Caffo, M. A. Lindquist, Comparing test-retest reliability of dynamic functional connectivity methods. Neuroimage. 158, 155–175 (2017).

65. E. Glerean, J. Salmi, J. M. Lahnakoski, I. P. Jääskeläinen, M. Sams, Functional magnetic resonance imaging phase synchronization as a measure of dynamic functional connectivity. Brain Connect. 2, 91–101 (2012).

66. C. Baldassano, J. Chen, A. Zadbood, J. W. Pillow, U. Hasson, K. A. Norman, Discovering Event Structure in Continuous Narrative Perception and Memory. Neuron. 95, 709–721.e5 (2017).

67. G. J. Stephens, L. J. Silbert, U. Hasson, Speaker–listener neural coupling underlies successful communication. Proceedings of the National Academy of Sciences. 107, 14425–14430 (2010).

68. H. W. Kuhn, The Hungarian method for the assignment problem. Naval research logistics quarterly (1955).

69. J. N. van der Meer, M. Breakspear, L. J. Chang, S. Sonkusare, L. Cocchi, Movie viewing elicits rich and reliable brain state dynamics. Nat. Commun. 11, 5004 (2020).

70. G. Schwarz, Estimating the Dimension of a Model. Ann. Stat. 6, 461–464 (1978).

71. Cheong, J.H., Brooks, S., & Chang, L.J., FaceSync: Open source framework for recording facial expressions with head-mounted cameras. F1000Research. 8 (2019) (available at https://f1000research.com/articles/8-702/v1).

72. P. Ekman, W. V. Friesen, Measuring facial movement. J. Nonverbal Behav. 1, 56–75 (1976).

73. G. Littlewort, J. Whitehill, T. Wu, I. Fasel, M. Frank, J. Movellan, M. Bartlett, in Face and Gesture 2011 (2011), pp. 298–305.

74. P.-H. (cameron) Chen, J. Chen, Y. Yeshurun, U. Hasson, J. Haxby, P. J. Ramadge, in Advances in Neural Information Processing Systems 28, C. Cortes, N. D. Lawrence, D. D. Lee, M. Sugiyama, R. Garnett, Eds. (Curran Associates, Inc., 2015), pp. 460–468.

75. A. C. Heusser, K. Ziman, L. L. W. Owen, HyperTools: a Python toolbox for gaining geometric insights into high-dimensional data. The Journal of Machine (2017) (available at https://dl.acm.org/doi/abs/10.5555/3122009.3242009).

76. J. F. Cohn, Z. Ambadar, P. Ekman, Observer-based measurement of facial expression with the Facial Action Coding System. The handbook of emotion elicitation and assessment, 203–221 (2007).

77. O. Kang, T. Wheatley, Pupil dilation patterns spontaneously synchronize across individuals during shared attention. J. Exp. Psychol. Gen. 146, 569–576 (2017).

78. L. J. Chang, J. Huckins, J. H. Cheong, S. Brietzke, M. A. Lindquist, T. D. Wager, ljchang/dartbrains: An online open access resource for learning functional neuroimaging analysis methods in Python (2020; https://zenodo.org/record/3909718).

79. L. J. Chang, T. Yarkoni, M. W. Khaw, A. G. Sanfey, Decoding the role of the insula in human cognition: functional parcellation and large-scale reverse inference. Cereb. Cortex. 23, 739–749 (2013).

80. S. Sul, B. Güroğlu, E. A. Crone, L. J. Chang, Medial prefrontal cortical thinning mediates shifts in other-regarding preferences during adolescence. Sci. Rep. 7, 8510 (2017).

81. C. F. Doeller, C. Barry, N. Burgess, Evidence for grid cells in a human memory network. Nature. 463, 657–661 (2010).

82. A. O. Constantinescu, J. X. O’Reilly, T. E. J. Behrens, Organizing conceptual knowledge in humans with a gridlike code. Science. 352, 1464–1468 (2016).

83. N. W. Schuck, M. B. Cai, R. C. Wilson, Y. Niv, Human orbitofrontal cortex represents a cognitive map of state space. Neuron. 91, 1402–1412 (2016).

84. G. J. Stephens, L. J. Silbert, Speaker–listener neural coupling underlies successful communication. Proceedings of the. 107, 14425–14430 (2010).

85. S. Suzuki, L. Cross, J. P. O’Doherty, Elucidating the underlying components of food valuation in the human orbitofrontal cortex. Nat. Neurosci. 20, 1780 (2017).

86. J. Reber, J. S. Feinstein, J. P. O’Doherty, M. Liljeholm, R. Adolphs, D. Tranel, Selective impairment of goal-directed decision-making following lesions to the human ventromedial prefrontal cortex. Brain. 140, 1743–1756 (2017).

87. M. J. F. Robinson, K. C. Berridge, Instant Transformation of Learned Repulsion into Motivational “Wanting.” Curr. Biol. 23, 282–289 (2013).

88. A. E. Skerry, R. Saxe, Neural representations of emotion are organized around abstract event features. Curr. Biol. 25, 1945–1954 (2015).

89. H. Saarimäki, L. F. Ejtehadian, E. Glerean, I. P. Jääskeläinen, P. Vuilleumier, M. Sams, L. Nummenmaa, Distributed affective space represents multiple emotion categories across the human brain. Soc. Cogn. Affect. Neurosci. 13, 471–482 (2018).

90. A. K. Anderson, K. Christoff, I. Stappen, D. Panitz, D. G. Ghahremani, G. Glover, J. D. E. Gabrieli, N. Sobel, Dissociated neural representations of intensity and valence in human olfaction. Nat. Neurosci. 6, 196–202 (2003).

91. J. A. Gottfried, R. Deichmann, J. S. Winston, R. J. Dolan, Functional heterogeneity in human olfactory cortex: an event-related functional magnetic resonance imaging study. J. Neurosci. 22, 10819–10828 (2002).

92. O. Lositsky, J. Chen, D. Toker, C. J. Honey, M. Shvartsman, J. L. Poppenk, U. Hasson, K. A. Norman, Neural pattern change during encoding of a narrative predicts retrospective duration estimates. Elife. 5, e16070 (2016).

93. A. C. Heusser, P. C. Fitzpatrick, J. R. Manning, How is experience transformed into memory? bioRxiv (2018), doi:10.1101/409987.

94. K. J. Friston, A. P. Holmes, K. J. Worsley, Statistical parametric maps in functional imaging: a general linear approach. Hum. Brain Mapp. (1994) (available at https://onlinelibrary.wiley.com/doi/abs/10.1002/hbm.460020402).

95. M. D. Fox, M. E. Raichle, Spontaneous fluctuations in brain activity observed with functional magnetic resonance imaging. Nat. Rev. Neurosci. 8, 700–711 (2007).

96. N. Kriegeskorte, R. Goebel, P. Bandettini, Information-based functional brain mapping. Proc. Natl. Acad. Sci. U. S. A. 103, 3863–3868 (2006).

97. J.-D. Haynes, G. Rees, Neuroimaging: decoding mental states from brain activity in humans. Nat. Rev. Neurosci. 7, 523 (2006).

98. R. Mukamel, H. Gelbard, A. Arieli, U. Hasson, I. Fried, R. Malach, Coupling between neuronal firing, field potentials, and FMRI in human auditory cortex. Science. 309, 951–954 (2005).

99. A. S. Heller, R. C. Lapate, K. E. Mayer, R. J. Davidson, The face of negative affect: trial-by-trial corrugator responses to negative pictures are positively associated with amygdala and negatively associated with ventromedial prefrontal cortex activity. J. Cogn. Neurosci. 26, 2102–2110 (2014).

100. L. J. Chang, P. J. Gianaros, S. B. Manuck, A. Krishnan, T. D. Wager, A Sensitive and Specific Neural Signature for Picture-Induced Negative Affect. PLoS Biol. 13, e1002180 (2015).

101. A. Krishnan, C.-W. Woo, L. J. Chang, L. Ruzic, X. Gu, M. López-Solà, P. L. Jackson, J. Pujol, J. Fan, T. D. Wager, Somatic and vicarious pain are represented by dissociable multivariate brain patterns. Elife. 5, e15166 (2016).

102. C.-W. Woo, L. J. Chang, M. A. Lindquist, T. D. Wager, Building better biomarkers: brain models in translational neuroimaging. Nat. Neurosci. 20, 365–377 (2017).

103. H. Eisenbarth, L. J. Chang, T. D. Wager, Multivariate Brain Prediction of Heart Rate and Skin Conductance Responses to Social Threat. J. Neurosci. 36, 11987–11998 (2016).

104. K. J. Gorgolewski, T. Auer, V. D. Calhoun, R. C. Craddock, S. Das, E. P. Duff, G. Flandin, S. S. Ghosh, T. Glatard, Y. O. Halchenko, D. A. Handwerker, M. Hanke, D. Keator, X. Li, Z. Michael, C. Maumet, B. N. Nichols, T. E. Nichols, J. Pellman, J.-B. Poline, A. Rokem, G. Schaefer, V. Sochat, W. Triplett, J. A. Turner, G. Varoquaux, R. A. Poldrack, The brain imaging data structure, a format for organizing and describing outputs of neuroimaging experiments. Sci Data. 3, 160044 (2016).

105. M. Visconti di Oleggio Castello, J. E. Dobson, T. Sackett, C. Kodiweera, J. V. Haxby, M. Goncalves, S. Ghosh, Y. O. Halchenko, ReproNim/reproin: 0.1.1 (http://dx.doi.org/10.5281/zenodo.1207118).

106. K. J. Gorgolewski, C. D. Burns, C. Madison, D. Clark, Y. O. Halchenko, M. L. Waskom, S. S. Ghosh, Nipype: a flexible, lightweight and extensible neuroimaging data processing framework in python. Front. Neuroinform. 5, 13 (2011).

107. K. J. Millman, M. Brett, Analysis of functional magnetic resonance imaging in python. Comput. Sci. Eng. 9, 52–55 (2007).

108. M. Jenkinson, P. Bannister, M. Brady, S. Smith, Improved optimization for the robust and accurate linear registration and motion correction of brain images. Neuroimage. 17, 825–841 (2002).

109. B. B. Avants, N. Tustison, G. Song, Advanced normalization tools (ANTS). Insight J. 2, 1–35 (2009).

110. M. Jenkinson, C. F. Beckmann, T. E. J. Behrens, M. W. Woolrich, S. M. Smith, FSL. Neuroimage. 62, 782–790 (2012).

111. S. Whitfield-Gabrieli, Artifact detection and QA manual. Massachusetts: Massachusetts Institute of Technology (2009).

112. L. Chang, J. Manning, C. Baldassano, A. de la Vega, G. Fleetwood, L. Geerligs, J. Haxby, J. Lahnakoski, C. Parkinson, H. Shappell, W. M. Shim, T. Wager, T. Yarkoni, Y. Yeshurun, E. Finn, naturalistic-data-analysis/naturalistic_data_analysis: Version 1.0 (2020; https://zenodo.org/record/3937849).

113. G. Chen, Y.-W. Shin, P. A. Taylor, D. R. Glen, R. C. Reynolds, R. B. Israel, R. W. Cox, Untangling the relatedness among correlations, part I: Nonparametric approaches to inter-subject correlation analysis at the group level. Neuroimage. 142, 248–259 (2016).

114. G. J. Székely, M. L. Rizzo, The Energy of Data. Annu. Rev. Stat. Appl. 4, 447–479 (2017).

115. G. J. Székely, M. L. Rizzo, N. K. Bakirov, Measuring and testing dependence by correlation of distances. Ann. Stat. 35, 2769–2794 (2007).

116. E. Jones, T. Oliphant, P. Peterson, {SciPy}: Open source scientific tools for {Python} (2001--), (available at http://www.scipy.org).

117. L. R. Rabiner, A tutorial on hidden Markov models and selected applications in speech recognition. Proc. IEEE. 77, 257–286 (1989).

118. J. A. Bilmes, Others, A gentle tutorial of the EM algorithm and its application to parameter estimation for Gaussian mixture and hidden Markov models. International Computer Science Institute. 4, 126 (1998).

119. T. Yarkoni, R. A. Poldrack, T. E. Nichols, D. C. Van Essen, T. D. Wager, Large-scale automated synthesis of human functional neuroimaging data. Nat. Methods. 8, 665–670 (2011).

120. A. S. Fox, L. J. Chang, K. J. Gorgolewski, T. Yarkoni, Bridging psychology and genetics using large-scale spatial analysis of neuroimaging and neurogenetic data. bioRxiv (2014), p. 012310.

121. P.-H. A. Chen, E. Jolly, J. H. Cheong, L. J. Chang, Intersubject representational similarity analysis reveals individual variations in affective experience when watching erotic movies. Neuroimage. 216, 116851 (2020).

122. iMotions Biometric Research Platform 6.0 (iMotions A/S, Copenhagen, Denmark, 2016).

123. J. H. Cheong, S. Byrnes, L. J. Chang, Facial expression analysis toolbox.

124. F. Pedregosa, G. Varoquaux, A. Gramfort, V. Michel, B. Thirion, O. Grisel, M. Blondel, P. Prettenhofer, R. Weiss, V. Dubourg, J. Vanderplas, A. Passos, D. Cournapeau, M. Brucher, M. Perrot, É. Duchesnay, Scikit-learn: machine learning in python. J. Mach. Learn. Res. 12, 2825–2830 (2011).

125. P. Lucey, J. F. Cohn, T. Kanade, J. Saragih, Z. Ambadar, I. Matthews, in 2010 IEEE Computer Society Conference on Computer Vision and Pattern Recognition - Workshops (ieeexplore.ieee.org, 2010), pp. 94–101.

126. T. Baltrušaitis, P. Robinson, L. P. Morency, in 2016 IEEE Winter Conference on Applications of Computer Vision (WACV) (ieeexplore.ieee.org, 2016), pp. 1–10.

127. L. J. Chang, N. Greenstein, H. Eisenbarth, M. Reddan, E. Andrews, T. D. Wager, Recovering individual sparse emotion ratings using collaborative filtering techniques.

128. G. Linden, B. Smith, J. York, Amazon.com recommendations: item-to-item collaborative filtering. IEEE Internet Comput. 7, 76–80 (2003).

129. X. Su, T. M. Khoshgoftaar, A survey of collaborative filtering techniques. Advances in Artificial Intelligence. 2009, 2 (2009).

130. L. Chang, E. Jolly, J. H. Cheong, A. Burnashev, A. Chen, cosanlab/nitools: 0.3.11 (2018; https://zenodo.org/record/2229813).

131. M. Brett, M. Hanke, B. Cipollini, M.-A. Côté, C. Markiewicz, S. Gerhard, E. Larson, G. R. Lee, Y. Halchenko, E. Kastman, Others, nibabel: 2.1. 0. Zenodo (2016).

132. A. Abraham, F. Pedregosa, M. Eickenberg, P. Gervais, A. Mueller, J. Kossaifi, A. Gramfort, B. Thirion, G. Varoquaux, Machine learning for neuroimaging with scikit-learn. Front. Neuroinform. 8, 14 (2014).

133. T. E. Oliphant, A guide to NumPy (Trelgol Publishing USA, 2006), vol. 1.

134. W. McKinney, Others, in Proceedings of the 9th Python in Science Conference (Austin, TX, 2010), vol. 445, pp. 51–56.

135. J. D. Hunter, Matplotlib: A 2D Graphics Environment. Computing in Science Engineering. 9, 90–95 (2007).

136. M. Waskom, O. Botvinnik, P. Hobson, J. Warmenhoven, Seaborn: statistical data visualization (2014).

