## Supplemental Materials for "Endogenous variation in ventromedial prefrontal cortex state dynamics during naturalistic viewing reflects affective experience"

#### Table of Contents

Supplemental Results

- *Voxel Autocorrelation*
- *Study 1 HMM Cross-Episode State Prediction*
- *Group HMM State Concordance Feature Mapping*

Supplemental Tables

- *Table S1. Similarity of Individual HMM states Study 1*
- *Table S2. Similarity of Individual HMM states Study 2*

Supplemental Figures

- *Figure S1. Temporal recurrence of spatial patterns in vmPFC plotted for each participant in Study 1*
- *Figure S2. Temporal recurrence of spatial patterns in V1 plotted for each participant in Study 1*
- *Figure S3. Temporal recurrence of spatial patterns in vmPFC plotted for each participant in Study 2*
- *Figure S4. Temporal recurrence of spatial patterns in V1 plotted for each participant in Study 2*
- *Figure S5. Voxelwise Autocorrelation*
- *Figure S6. Study 1 vmPFC HMM Spatial Patterns*
- *Figure S7. Study 2 vmPFC HMM Spatial Patterns*
- *Figure S8. Cross Episode Group HMM State Concordance*
- *Figure S9. Group HMM vmPFC State Transition Probabilities*
- *Figure S10. Group HMM Model Fits*
- *Figure S11. PCC Analyses*
- *Figure S12. V1 Group-HMM state concordance and selected features*
- *Figure S13. PCC Group-HMM state concordance and selected features*
- *Figure S14. vmPFC Group-HMM state concordance and selected features*
- *Figure S15. Visual and affective features associated with high state concordance*
- *Figure S16. Inter-Experiment Shared Response Model Face Expression Weights*
- *Figure S17. Inter-Experiment Shared Response Model Emotion Rating Weights*
- *Figure S18. HMM State Contrast Decoding.*

### Supplemental Results

#### Voxel Autocorrelation

To further explore temporal autocorrelations during naturalistic viewing, we estimated the autocorrelation, up to 50 lagged TRs, for each voxel in the brain for each participant across Study 1 and Study 2. We modeled the decay using a three parameter exponential function (see methods). For each voxel, we computed the time (in TRs) it took for the predicted autocorrelation function to decay to a value of 0.1. This yielded a single decay duration for each voxel. Overall, these per-voxel autocorrelation decay durations were highly consistent across scanning sessions. We correlated the average voxel-wise autocorrelation maps (of the decay durations) for participants in Study 1 that were scanned watching episodes 1 and 2, and found a high level of reliability across the whole brain ($r = 0.91$, $p < 0.001$) and within the vmPFC ($r = 0.88$, $p < 0.001$). The reliability across scanners (i.e., Study 1 Phillips Achieva Intera and Study 2 Siemens Prisma) was considerably lower across the whole brain ($r = 0.72$, $p < 0.001$) and within the vmPFC ($r = 0.55$, $p < 0.001$). As can be seen in Figure S5, the voxels on the ventral surface of the orbital frontal cortex exhibited the longest temporal autocorrelation. These voxels are also presumably the most likely to be impacted by echo planar imaging susceptibility artifacts. To explore this relationship, we computed the average temporal signal to noise ratio (tSNR) for each voxel on the preprocessed data (not denoised) and correlated the average autocorrelation duration with the average tSNR for each voxel within the vmPFC (Figure S5C). We observed a strong relationship between voxel autocorrelation and tSNR for Study 1 ($r = -0.72$, $p < 0.001$) and Study 2 ($r = -0.71$, $p < 0.001$). This highlights an important limitation in interpreting our autocorrelation findings at the voxel level. To assess the impact of susceptibility artifacts on our results, we regressed out average tSNR from each voxel for each study within the vmPFC and re-computed the reliability of the autocorrelation duration across studies. This resulted in a numeric *increase* in reliability ($r = 0.58$, $p < 0.001$; Figure S5D). Though we think it is important to interpret our voxelwise autocorrelation results with caution, we do not believe that our observed effects can be solely attributed to signal dropout or spatial distortions resulting from susceptibility artifacts.

#### Study 1 HMM Cross-Episode State Prediction

In the main text, we reported analyses demonstrating the reliability of the Group HMM model. Broadly, we found that the Group HMM models identify consistent states across two independent samples when watching episode 1 of Friday Night Lights. Our inter-experiment latent factor model revealed that some of these states appeared to reflect latent emotional states that were all manifested in facial expressions, and ratings of subjective feelings from two additional independent samples of participants. We were additionally interested in the generalizability of the states across other contexts. To assess the generalizability, we focused specifically on participants from Study1 and trained an additional k=4 Group HMM on the PCA reduced vmPFC data from viewing episode 2. We aligned the states from the models trained on each episode using the Hungarian algorithm (Kuhn 1955). We then assessed the cross-episode state concordance synchronization using Pearson correlations. Overall, we found that three of the four states appeared to be capturing similar states when testing the models on episode 1 (state1: $r=0.3$, $p < 0.001$; state 2: $r=0.64$, $p < 0.001$; state 3: $r=1.0$, $p = 0.001$; state 4: $r=-0.03$, $p = 0.29$; Figure S8A) and episode 2 (state1: $r=0.3$, $p = 003$; state 2: $r=0.71$, $p = 0.002$; state 3: $r=1.0$, $p < 0.001$; state 4: $r=0.01$, $p = 0.93$; Figure S8B) using a circle-shifting permutation approach with 5,000 random shifts. The full temporal state concordance correlation matrix averaged across episodes is displayed in Figure S8C. In addition, three out of the four states shared similar voxelwise spatial patterns across the two episodes (state1: $r=0.28$, $p < 0.001$; state 2: $r=0.28$, $p < 0.001$; state 3: $r=0.03$, $p = 0.11$; state 4: $r=0.11$, $p < 0.001$; Figure S8D). These results suggest that the vmPFC states estimated while viewing a single episode may generalize to other episodes. State 3 is likely capturing some type of unexplained variability or "noise," as the spatial patterns are not spatially similar and we found consistently low concordance in State 3 across both episodes.

#### Group HMM State Concordance Feature Mapping

Our inter-experiment shared response model identified facial expression and subjective feelings that displayed similar temporal dynamics as the participant state concordance fluctuated throughout the viewing episode. When more participants from Studies 1 & 2 were occupying the same vmPFC state, participants from Studies 3 & 4 were more likely to be in a positive or negative affective state. Because the affective states were elicited by the television show, it is not possible to separate feelings from other perceptual processes such as visual or auditory processing. For example, the music might change during scenes that are sad or more celebratory. We ran several additional analyses to explore the specificity of these affective processes to several other regions in addition to the vmPFC including V1 and PCC.

First, we have visualized the temporal dynamics of several different visual and affective features with state concordances from Group-HMMs fit separately fit to each ROI (i.e., V1, PCC, and vmPFC) (Figure S12, S13, & S14). These features include (a) visual brightness based on the average luminosity of pixels from each frame from Friday Night Lights episode 1, (b) visual vibrance based on the variance of the color channels for each frame from Friday Night Lights episode 1, (c) average AU12 Lip Corner Puller intensity (i.e., smiling) from Study 3, (d) average AU15 Lip Corner Depressor (e.g., frowning) from Study 3, (e) average subjective Joy ratings from Study 4, and (f) average subjective Sadness ratings from Study 4. We highlight timepoints in which the state concordance exceeds an arbitrary threshold of 0.7. States that never reach a maximum concordance of at least 0.7 are excluded from this analysis. All features are normalized using the z-transform, except for Emotion Ratings, which are centered and then stacked to compute a Global Z-Score to maintain between rating differences, then convolved with a double gamma HRF (Glover 1999). Overall, we find that V1 state concordance appears to closely follow large changes in screen brightness. Consistent with the inter-experiment shared response model analysis, vmPFC State 1 tracks large changes in Joy ratings and smiling (AU12), while vmPFC State 2 is more strongly associated with sadness ratings and frowning (AU15) (see Figure S14). PCC appears to track large segments in which there is a large shift in visual luminance, but not large changes in any of the affective features. These results provide additional qualitative support to our interpretation that V1 state changes track visual processing, while vmPFC state changes appear to reflect changes in affective processing.

Second, we attempted to more quantitatively map the intensity of each feature to ROI state concordance fluctuations. Similar to the previous analyses described above, for each state in Studies 1 & 2, we extracted the average convolved feature intensity when at least 70% of the participants occupied a particular state and compared that to the average feature intensity at all other time points. We compared this contrast of high vs low state concordance to a null distribution of contrasts generated by repeatedly and randomly circularly shifting the feature time course. This procedure provides a non-parametric permutation test that preserves any temporal structure present in the data (i.e., autocorrelations, periodicities, etc.). For plotting purposes, we took the absolute value of each contrast and averaged all states from each study to separately illustrate the average signal intensity changes related to visual and affective processes in each ROI (Figures S15). We separately plot features related to visual processes including video luminance and action units 43 (Eyes Closed) & 5 (Upper Lid Raiser) that are related to opening and closing eyes, and affective features including subjective feeling ratings and action units measured in Study 3 & 4. This analysis provides additional quantitative support that V1 state changes are more associated with changes in features related to visual processing (i.e., luminance and closing eyes), while vmPFC is more associated with changes in affect - particularly sadness ratings and inner brow raiser (AU1) furrowing brow (AU4). PCC appears to also reflect changes in visual and affective processing including both sadness, guilt, satisfaction, and shame ratings and smiling (AU12 & AU14).

### Supplemental Tables

**Table S1.** Similarity of Individual HMM states Study 1

| ROI | State | Spatial Mean | Concordance Mean | Concordance Std |
|---|---|---|---|---|
| vmPFC | 0 | 0.66 | 0.25 | 0.12 |
| | 1 | 0.23 | 0.28 | 0.13 |
| | 2 | 0.16 | 0.26 | 0.11 |
| | 3 | 0.25 | 0.21 | 0.11 |
| V1 | 0 | 0.08 | 0.14 | 0.09 |
| | 1 | 0.20 | 0.20 | 0.11 |
| | 2 | 0.06 | 0.18 | 0.10 |
| | 3 | 0.87 | 0.16 | 0.11 |
| | 4 | 0.49 | 0.18 | 0.11 |
| | 5 | 0.26 | 0.14 | 0.09 |
| PCC | 0 | 0.31 | 0.06 | 0.07 |
| | 1 | 0.25 | 0.06 | 0.06 |
| | 2 | 0.19 | 0.07 | 0.07 |
| | 3 | 0.39 | 0/06 | 0.06 |
| | 4 | 0.18 | 0.05 | 0.07 |
| | 5 | 0.18 | 0.06 | 0.06 |
| | 6 | 0.15 | 0.05 | 0.07 |
| | 7 | 0.29 | 0.05 | 0.06 |
| | 8 | 0.17 | 0.05 | 0.06 |
| | 9 | 0.19 | 0.05 | 0.06 |
| | 10 | 0.30 | 0.05 | 0.06 |
| | 11 | 0.11 | 0.06 | 0.07 |
| | 12 | 0.28 | 0.06 | 0.07 |
| | 13 | 0.17 | 0.06 | 0.06 |
| | 14 | 0.16 | 0.06 | 0.07 |
| | 15 | 0.17 | 0.07 | 0.07 |
| | 16 | 0.03 | 0.06 | 0.06 |

**Table S2.** Similarity of Individual HMM states Study 2

| ROI | State | Spatial Mean | Concordance Mean | Concordance Std |
|---|---|---|---|---|
| vmPFC | 0 | 0.18 | 0.25 | 0.06 |
| | 1 | 0.34 | 0.25 | 0.08 |
| | 2 | 0.23 | 0.26 | 0.08 |
| | 3 | 0.26 | 0.25 | 0.07 |
| V1 | 0 | 0.71 | 0.13 | 0.06 |
| | 1 | 0.01 | 0.16 | 0.06 |
| | 2 | 0.59 | 0.15 | 0.06 |
| | 3 | 0.15 | 0.17 | 0.06 |
| | 4 | 0.07 | 0.15 | 0.06 |
| | 5 | 0.13 | 0.13 | 0.05 |
| | 6 | 0.35 | 0.12 | 0.05 |
| PCC | 0 | 0.18 | 0.09 | 0.04 |
| | 1 | 0.33 | 0.10 | 0.05 |
| | 2 | 0.34 | 0.08 | 0.05 |
| | 3 | 0.06 | 0.09 | 0.05 |
| | 4 | 0.17 | 0.08 | 0.04 |
| | 5 | 0.34 | 0.08 | 0.05 |
| | 6 | 0.25 | 0.09 | 005 |
| | 7 | 0.20 | 0.07 | 0.04 |
| | 8 | 0.19 | 0.08 | 0.05 |
| | 9 | 0.04 | 0.07 | 0.04 |
| | 10 | 0.18 | 0.09 | 0.05 |
| | 11 | 0.16 | 0.08 | 0.05 |

#### Supplemental Figures

**Figure S1. Temporal recurrence of spatial patterns in vmPFC plotted for each participant in Study 1.** Each plot represents the similarity of vmPFC response patterns over time (45-minutes) for a single participant (Study 1). Spatial patterns are correlated over time such that each cell in the matrix represents the pattern similarity between two given TRs. Color scale ranges between –1 (anticorrelated; blue) and 1 (maximally correlated; red).

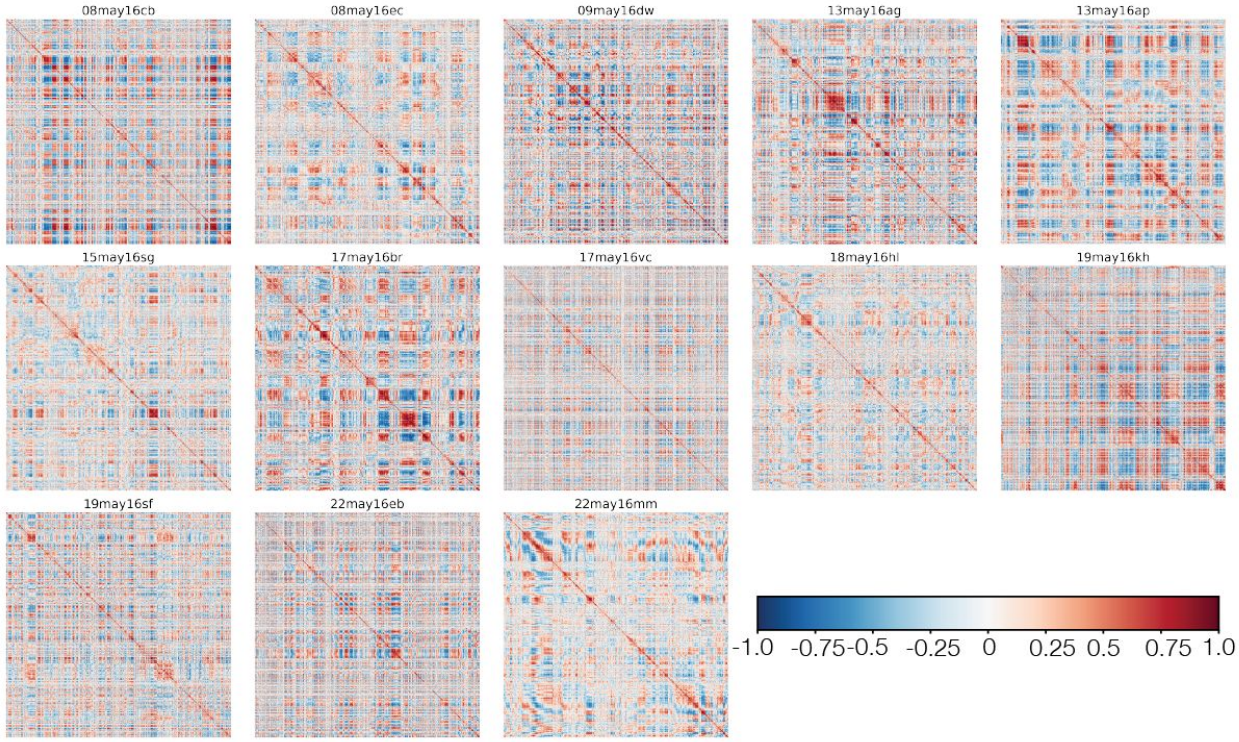

**Figure S2. Temporal recurrence of spatial patterns in V1 plotted for each participant in Study 1.** Each plot represents the similarity of V1 response patterns over time (45-minutes) for a single participant (Study 1). Spatial patterns are correlated over time such that each cell in the matrix represents the pattern similarity between two given TRs. Color scale ranges between –1 (anticorrelated; blue) and 1 (maximally correlated; red)

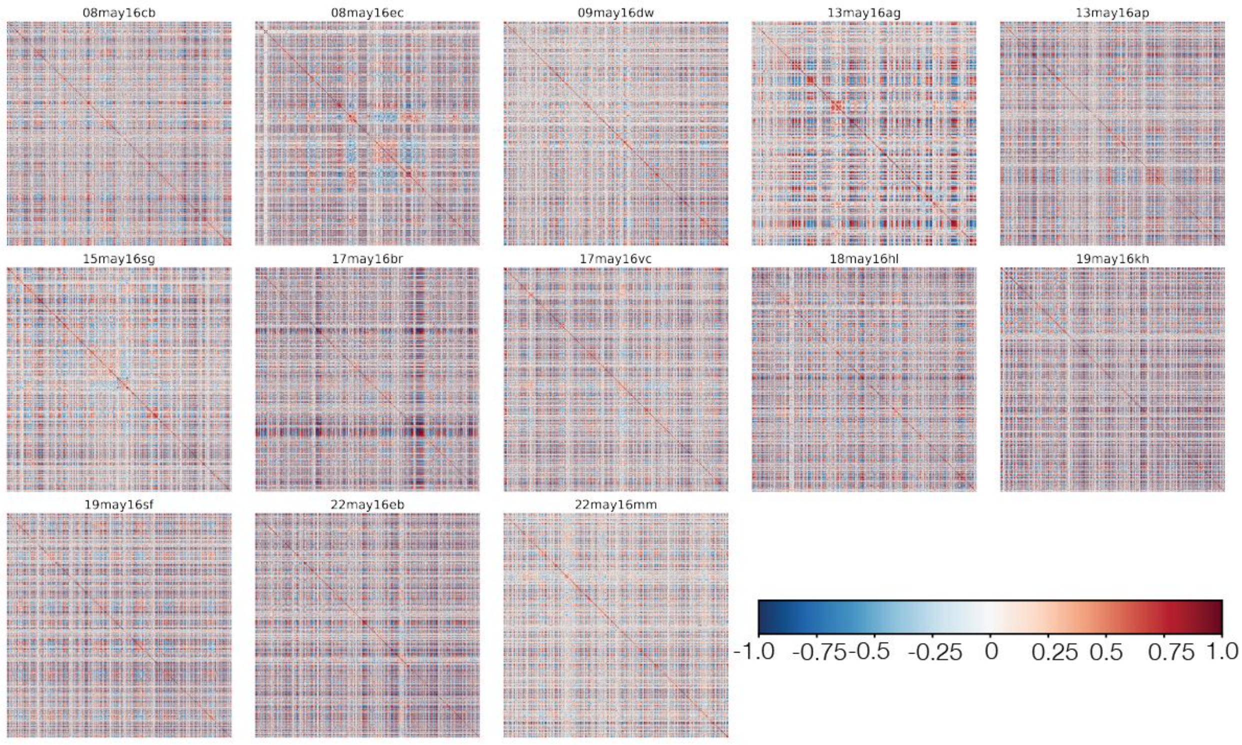

**Figure S3. Temporal recurrence of spatial patterns in vmPFC plotted for each participant in Study 2.** Each plot represents the similarity of vmPFC response patterns over time (45-minutes) for a single participant (Study 2). Spatial patterns are correlated over time such that each cell in the matrix represents the pattern similarity between two given TRs. Color scale ranges between –1 (anticorrelated; blue) and 1 (maximally correlated; red).

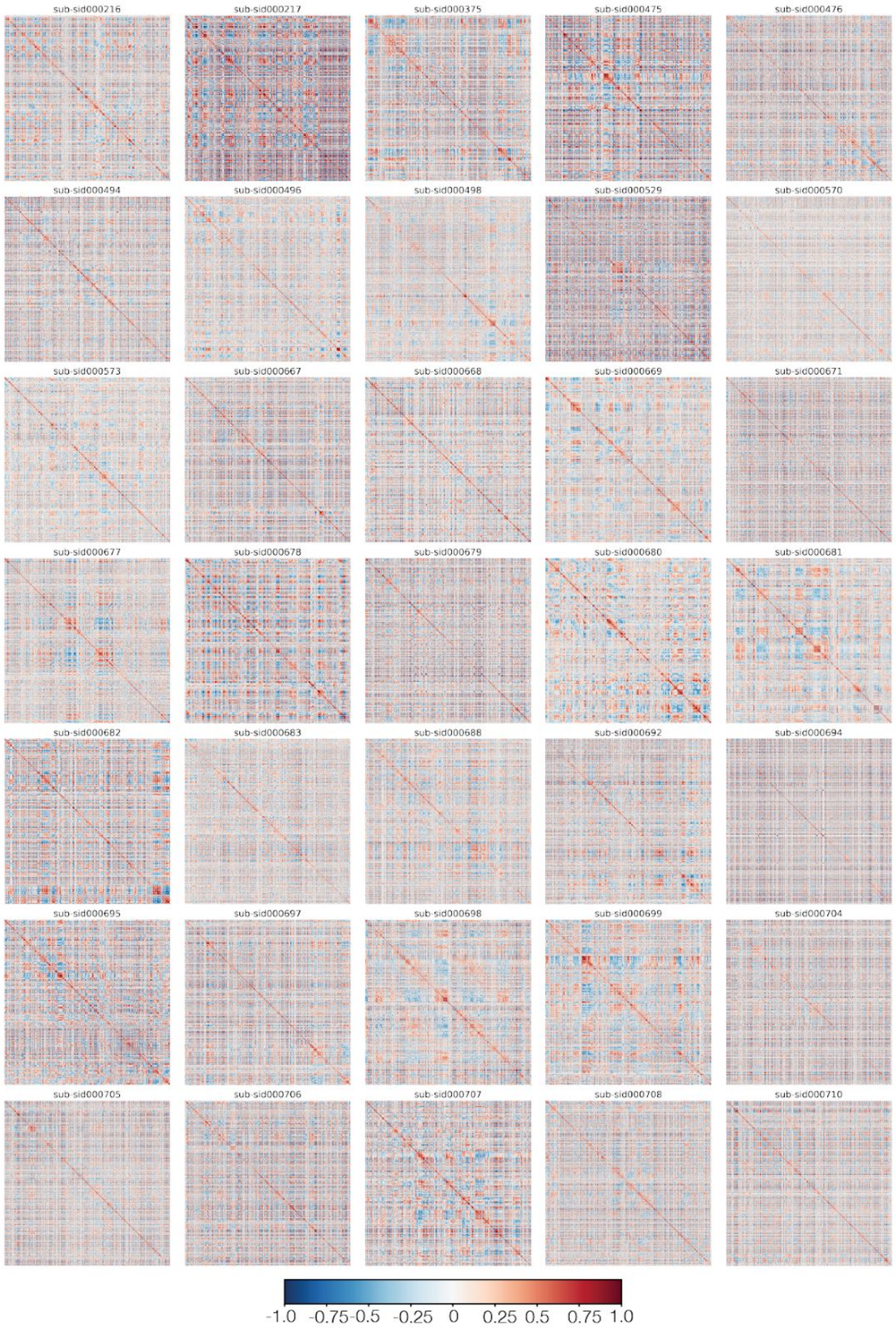

**Figure S4. Temporal recurrence of spatial patterns in V1 plotted for each participant in Study 2.** Each plot represents the similarity of V1 response patterns over time (45-minutes) for a single participant (Study 2). Spatial patterns are correlated over time such that each cell in the matrix represents the pattern similarity between two given TRs. Color scale ranges between –1 (anticorrelated; blue) and 1 (maximally correlated; red).

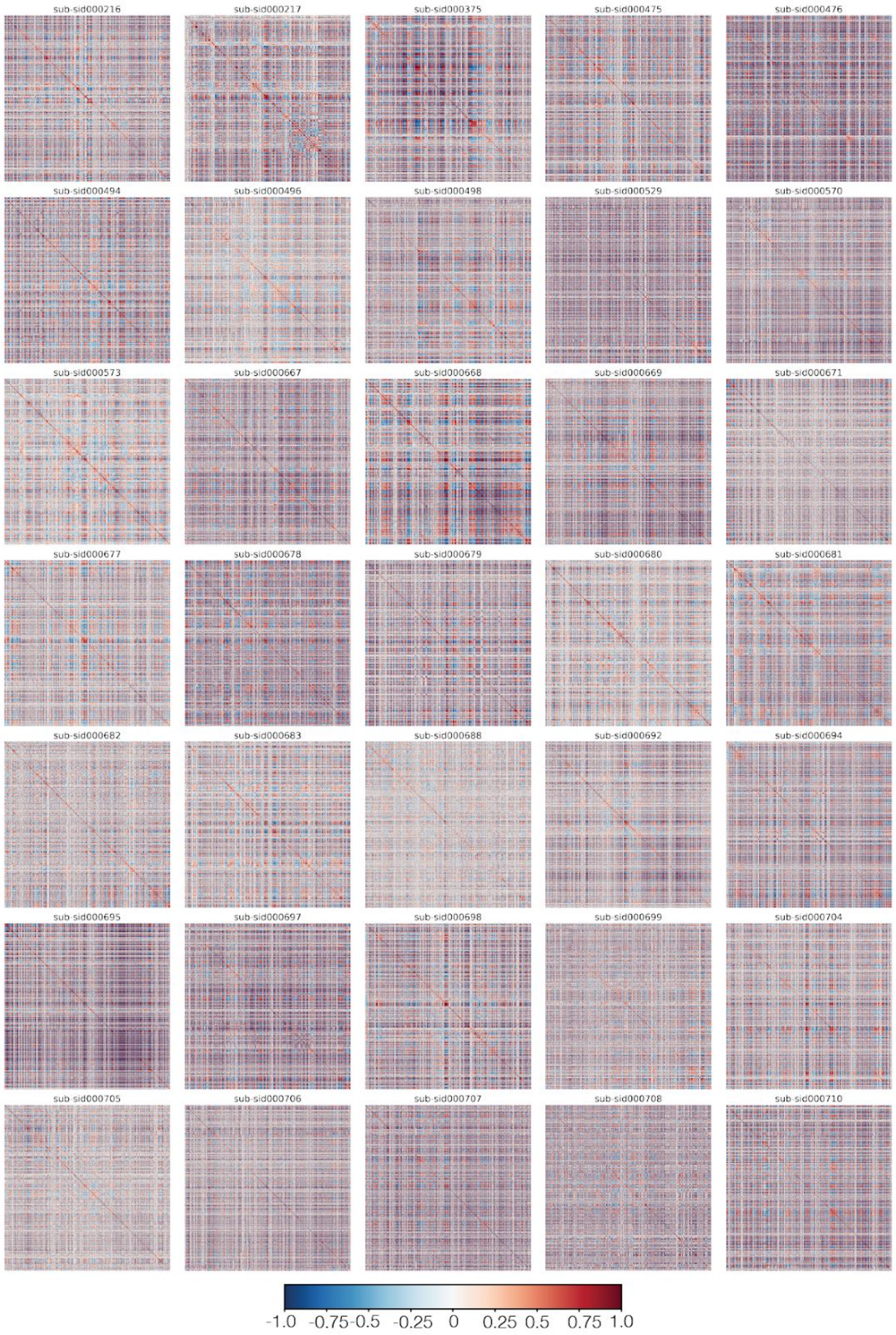

**Figure S5. Voxelwise Autocorrelation**. Panel's A & B depict the average time for the autocorrelation to decay to arbitrary 0.1 correlation for Study 1 and Study 2 respectively. In addition, we also plot the average voxelwise temporal signal to noise ratio (tSNR). Panel C illustrates the relationship between average tSNR and average autocorrelation duration across voxels for Study 1 and Study 2. Panel D shows the cross-study reliability of voxel autocorrelation duration within the vmPFC with and without removing variance attributed to tSNR.

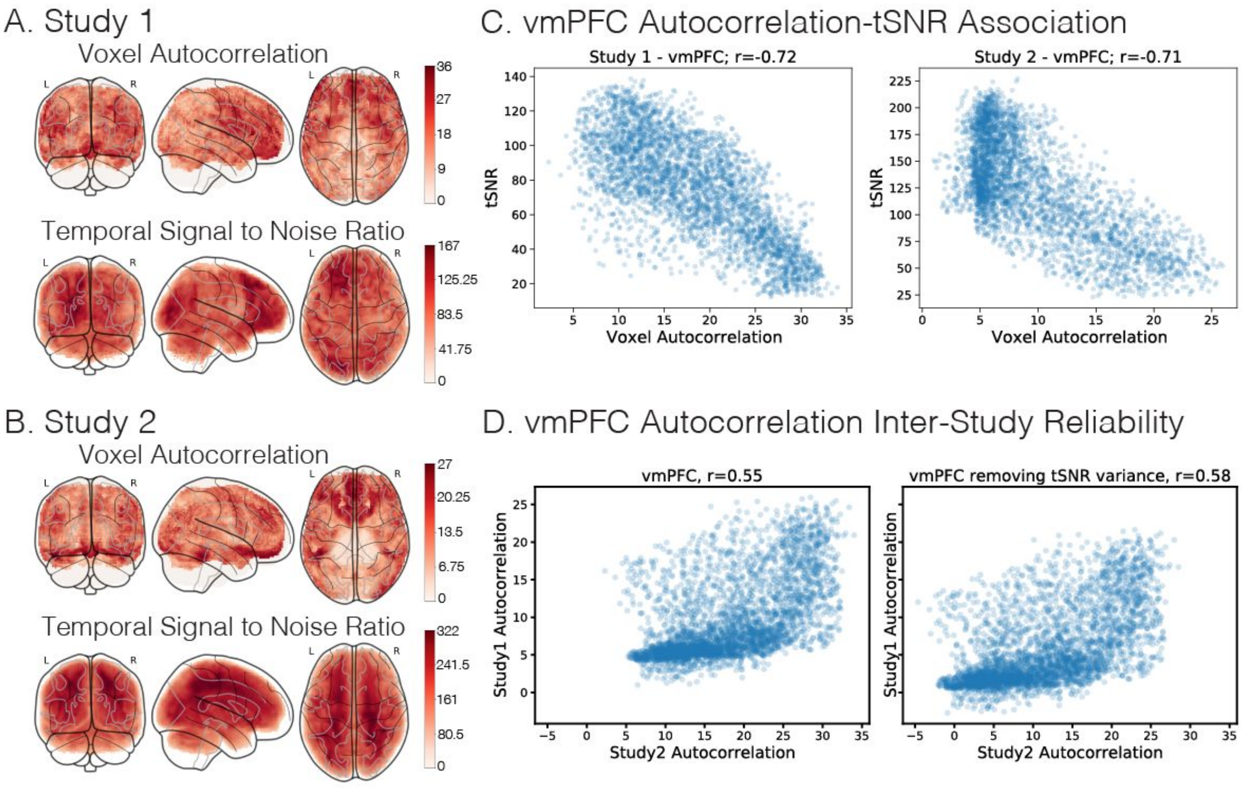

**Figure S6. Study 1 vmPFC HMM Spatial Patterns**. HMM Spatial emission patterns for each participant (n=13) from Study 1 (k=4). Each row is a separate participant. Each column is a distinct state.

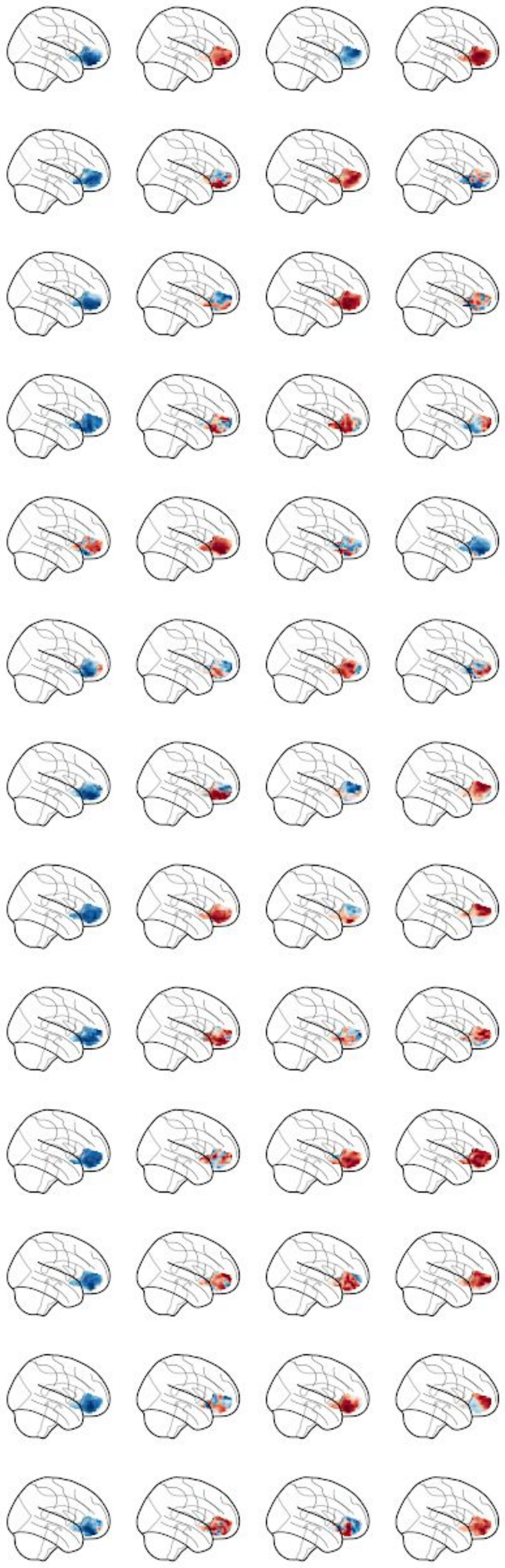

**Figure S7. Study 2 vmPFC HMM Spatial Patterns**. HMM Spatial emission patterns for each participant (n=35) from Study 2 (k=4). Each row is a separate participant. Each column is a distinct state.

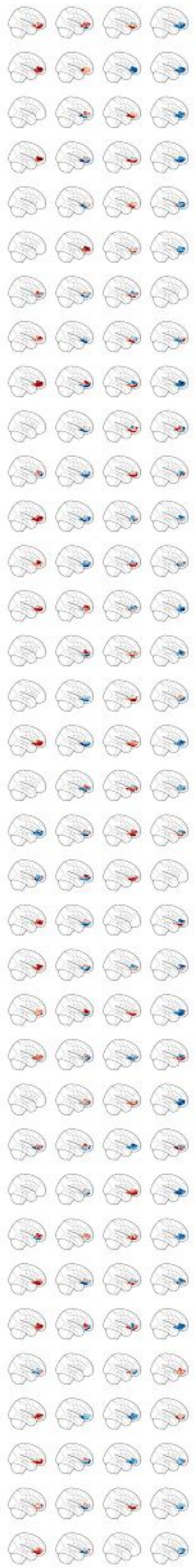

**Figure S8. Cross Episode Group HMM State Concordance**. Separate Group HMM k=4 Models were trained on data from Study 1 Episodes 1 and 2. Each model was used to decode the state sequences for each subject across both episodes using the Viterbi algorithm. State concordances for each model were aligned using the hungarian algorithm. A) Group HMM State concordance for Episode 1. B) Group HMM State concordance for Episode 2. C) Temporal similarity of state concordances averaged across episodes. D) Spatial similarity of state patterns. Red squares in Panel's C & D highlight the cross episode generalizability of each state.

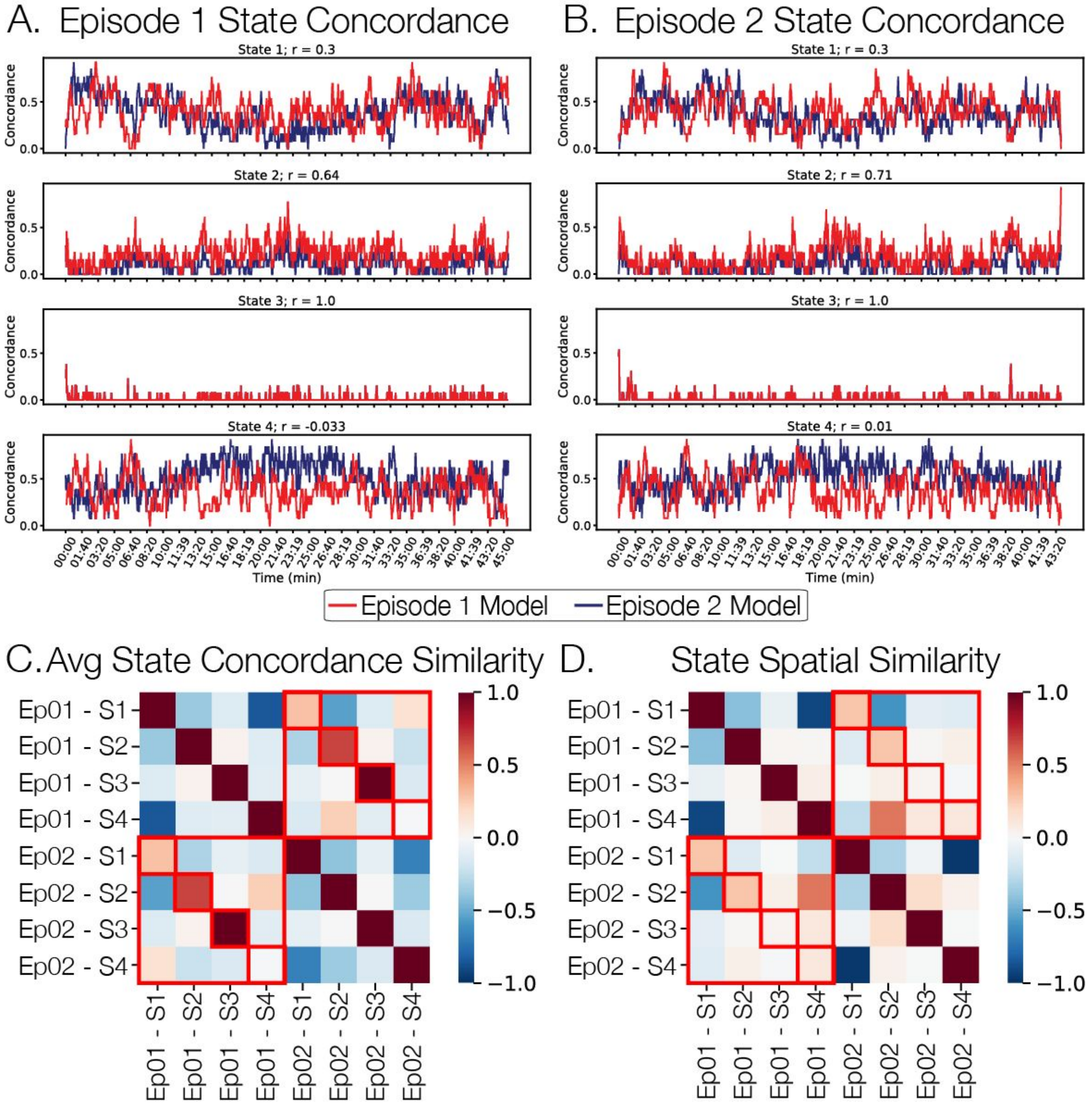

**Figure S9. Group HMM vmPFC State Transition Probabilities**. State transition probabilities estimated from study specific Group HMM k=4. A) Transition probabilities for each state estimated from Study 1 vmPFC activity. B) Transition probabilities for each state estimated from Study 2 vmPFC activity.

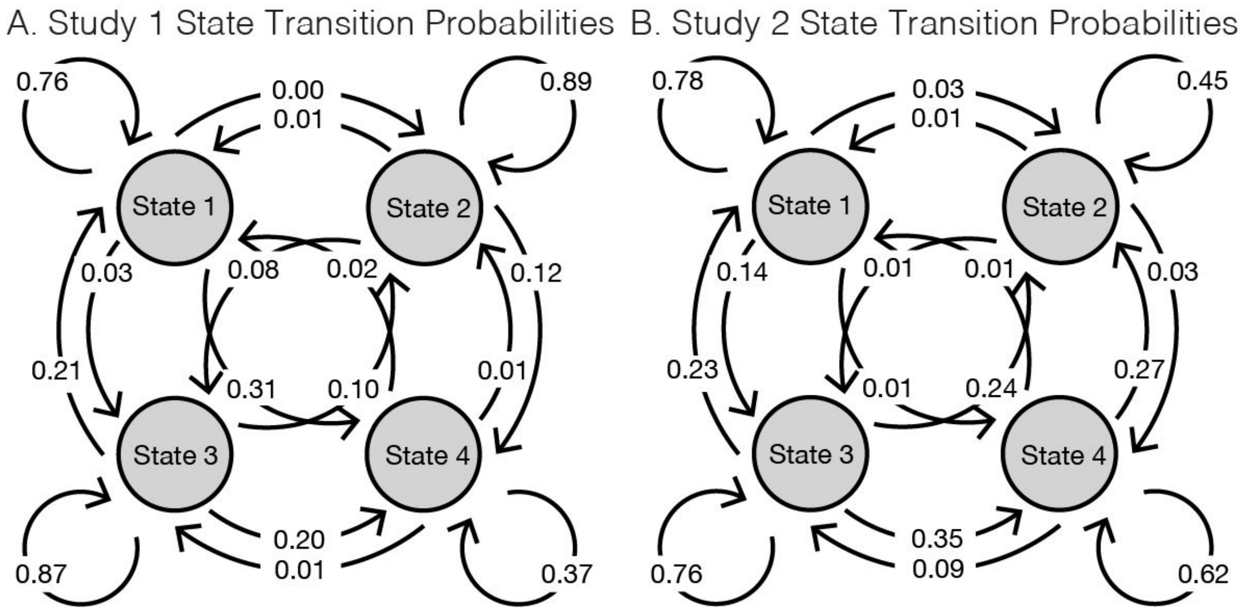

**Figure S10. Group HMM Model Fits**. Here we plot the Bayesian Information Criterion for states k=[2,25] for Group-HMM fit separately to V1, Posterior Cingulate Cortex (PCC), and Ventromedial Prefrontal Cortex (vmPFC) for Studies 1 & 2. BIC values are z-scored and red dotted lines indicate the largest drop in BIC with respect to the number of states (V1: k=3; PCC: k=3; vmPFC: k=4).

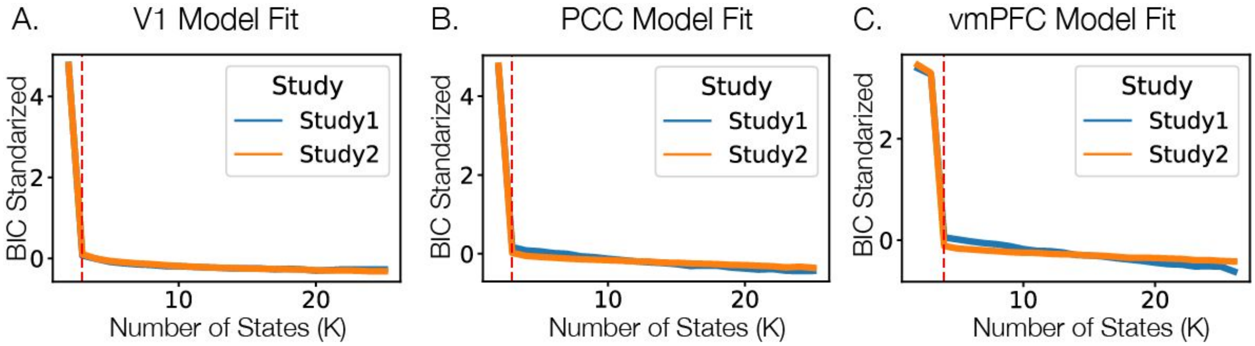

**Figure S11. PCC Analyses**. Here we plot all of the main analyses for an additional region of interest - the Posterior Cingulate Cortex (PCC). Panel A plots the average spatial ISC for Study 1 for both the anatomically and functionally aligned data. Panel B depicts the thresholded temporal recurrence matrix and subject by subject spatiotemporal similarity for Study 2, FDR q < 005 corresponds to p < 0.002. The average spatiotemporal similarity, r = 0.05, sd=0.03. Panel C depicts the autocorrelation at the voxel and pattern levels for both Study 1 and Study 2. Panel D depicts the results of the Individual-HMM analyses including (a) the spatial similarity of the k=17 patterns for Study 1 and k=12 patterns for Study 2, (b) the average spatial similarity for the cluster with the highest spatial similarity, (c) the average spatial patterns for each of the max spatial similarity clusters, and (d) the temporal state concordance of each of the max spatial similarity clusters.

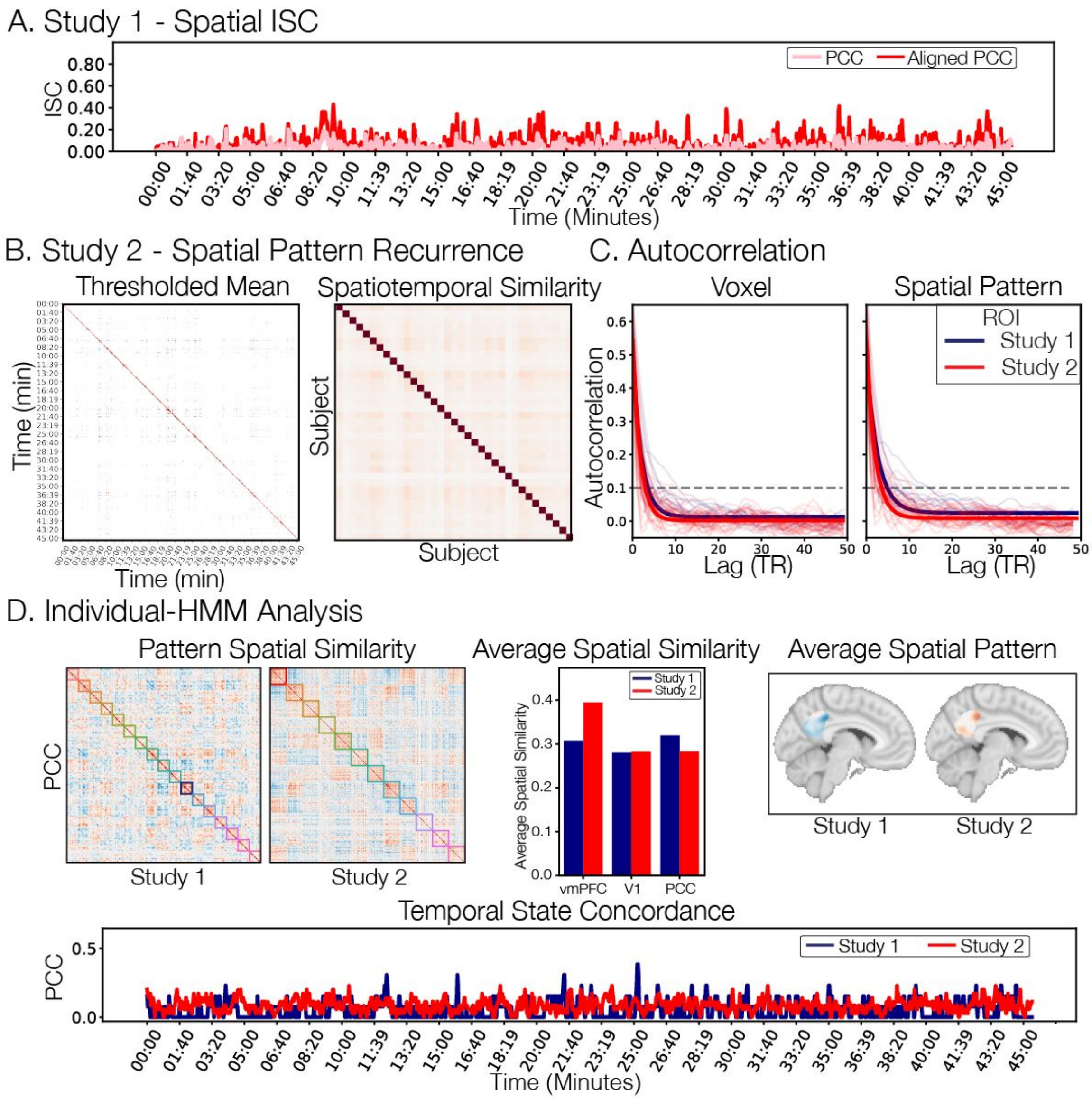

**Figure S12. V1 Group-HMM state concordance and selected features**. Here we plot the subject state concordance for V1 estimated from the Group-HMM k=3 along with several visual and affective features. We include: (a) visual brightness based on the average luminosity of pixels from each frame from Friday Night Lights episode 1, (b) visual vibrance based on the variance of the color channels for each frame from Friday Night Lights episode 1, (c) average AU12 Lip Corner Puller intensity (i.e., smiling) from Study 3, (d) average AU15 Lip Corner Depressor (e.g., frowning) from Study 3, (e) average subjective Joy ratings from Study 4, and (f) average subjective Sadness ratings from Study 4. All features are normalized using the z-transform, except for Emotion Ratings, which are centered and then stacked to compute a Global Z-Score to maintain between rating differences. The left column depicts V1 State 1 concordance and the right column depicts V1 State 2 concordance. Dotted red lines indicate any time point, in which the HMM State concordance from Study 1 exceeds 70% of the sample. V1 State 3 did not yield any time points that exceeded this threshold and therefore is not included.

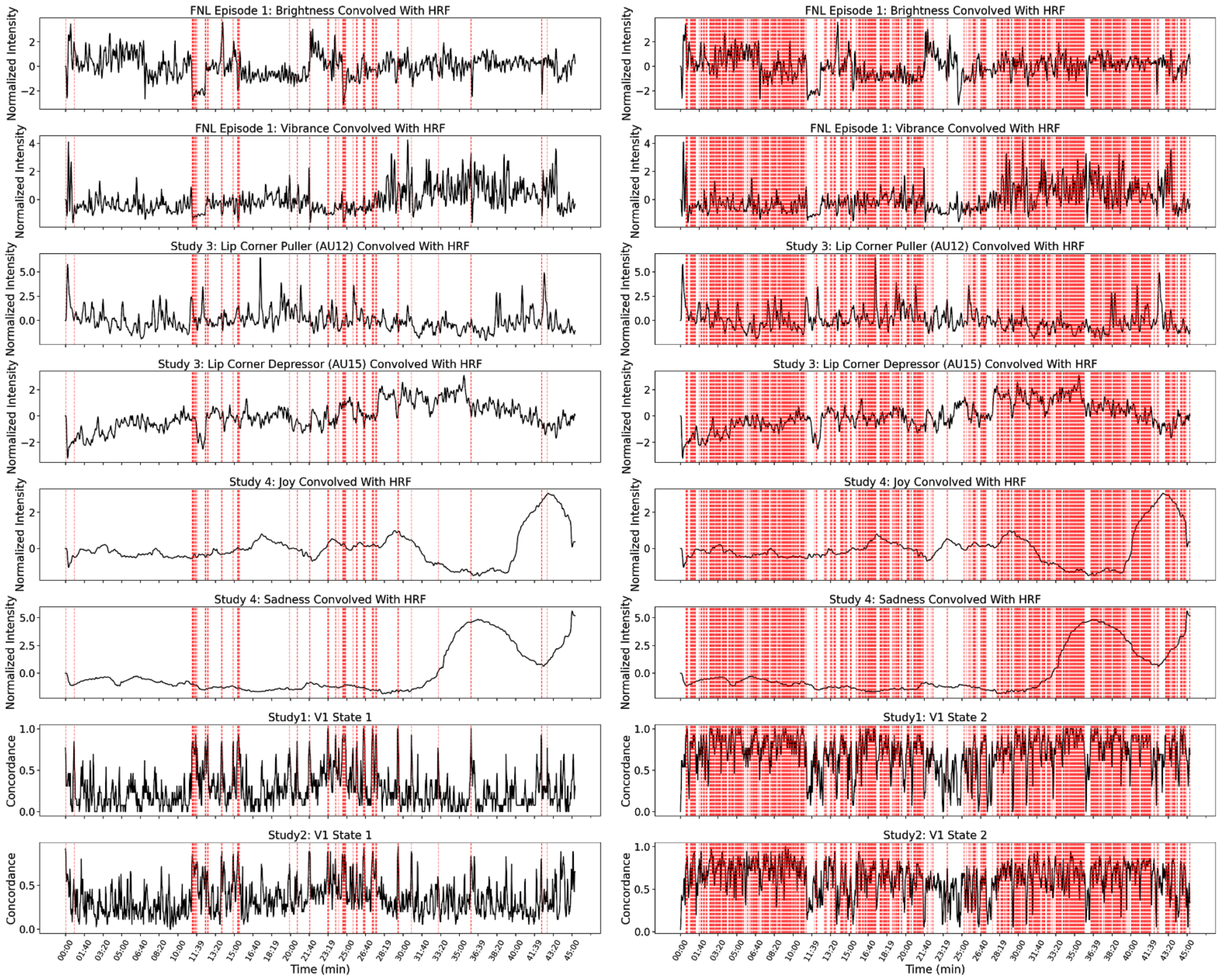

**Figure S13. PCC Group-HMM state concordance and selected features**. Here we plot the subject state concordance for PCC estimated from the Group-HMM k=3 along with several visual and affective features. We include: (a) visual brightness based on the average luminosity of pixels from each frame from Friday Night Lights episode 1, (b) visual vibrance based on the variance of the color channels for each frame from Friday Night Lights episode 1, (c) average AU12 Lip Corner Puller intensity (i.e., smiling) from Study 3, (d) average AU15 Lip Corner Depressor (e.g., frowning) from Study 3, (e) average subjective Joy ratings from Study 4, and (f) average subjective Sadness ratings from Study 4. All features are normalized using the z-transform, except for Emotion Ratings, which are centered and then stacked to compute a Global Z-Score to maintain between rating differences. The left column depicts PCC State 1 concordance and the right column depicts PCC State 2 concordance. Dotted red lines indicate any time point, in which the HMM State concordance from Study 1 exceeds 70% of the sample. PCC State 3 did not yield any time points that exceeded this threshold and therefore is not included.

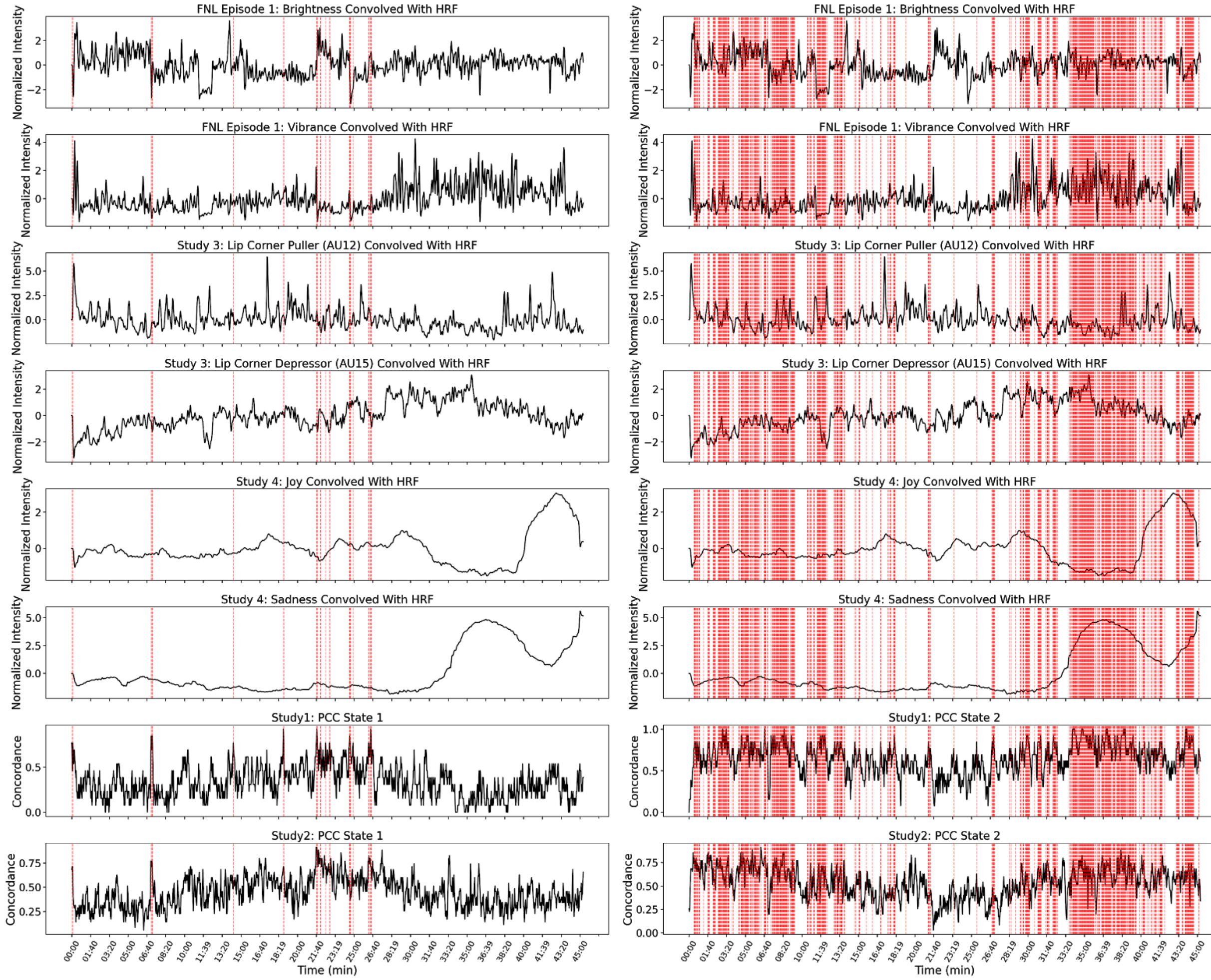

**Figure S14. vmPFC Group-HMM state concordance and selected features**. Here we plot the subject state concordance for vmPFC estimated from the Group-HMM k=3 along with several visual and affective features. We include: (a) visual brightness based on the average luminosity of pixels from each frame from Friday Night Lights episode 1, (b) visual vibrance based on the variance of the color channels for each frame from Friday Night Lights episode 1, (c) average AU12 Lip Corner Puller intensity (i.e., smiling) from Study 3, (d) average AU15 Lip Corner Depressor (e.g., frowning) from Study 3, (e) average subjective Joy ratings from Study 4, and (f) average subjective Sadness ratings from Study 4. All features are normalized using the z-transform, except for Emotion Ratings, which are centered and then stacked to compute a Global Z-Score to maintain between rating differences. The left column depicts vmPFC State 1 concordance and the right column depicts vmPFC State 2 concordance. Dotted red lines indicate any time point, in which the HMM State concordance from Study 1 exceeds 70% of the sample. vmPFC States 3 & 4 did not yield any time points that exceeded this threshold and therefore are not included.

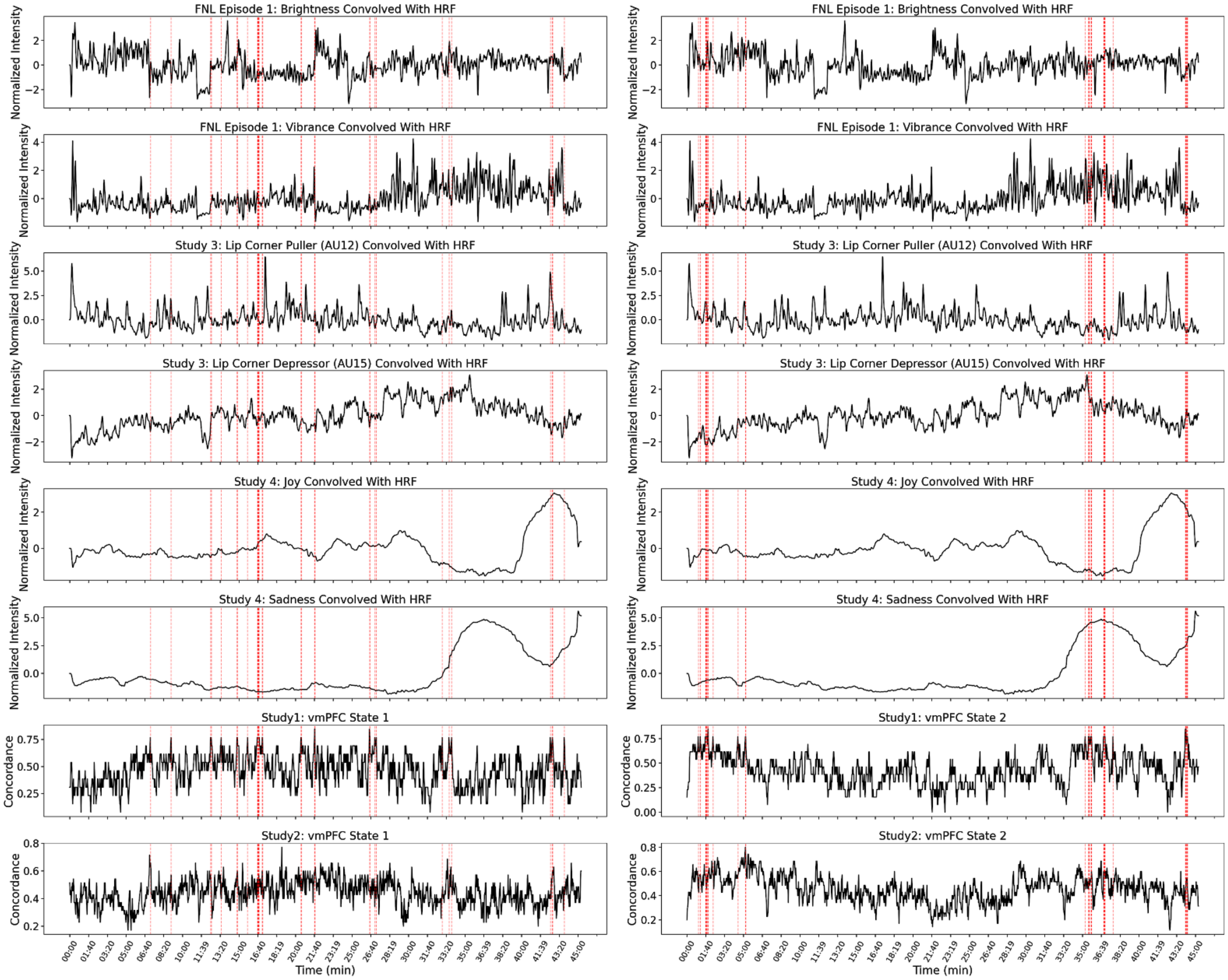

**Figure S15. Visual and affective features associated with high state concordance.** Here we highlight features convolved with a double-gamma HRF that were significantly different during time periods of high state concordance compared to low state concordance. For each fMRI study (Study 1 & 2), we computed the average feature intensity when the state concordance for each state identified from the Group-HMM exceeded 70% of the sample and compared it to the average feature intensity for all other time points. We took the absolute value of the mean difference and averaged across states and studies. We plotted features that were statistically significant ($p < 0.05$) based on a permutation test in which we separately generated a null distribution for each feature by randomly circularly shifting the feature time series. This permutation approach preserves any temporal structure inherently present in the data. We separately plotted features related to visual processes including video luminance and action units 43 (Eyes Closed) & 5 (Upper Lid Raiser) that are related to opening and closing eyes, from affective features including subjective feeling ratings and action units measured in Study 3 & 4. We have separated these based on Group-HMM states estimated from V1, PCC, and vmPFC regions.

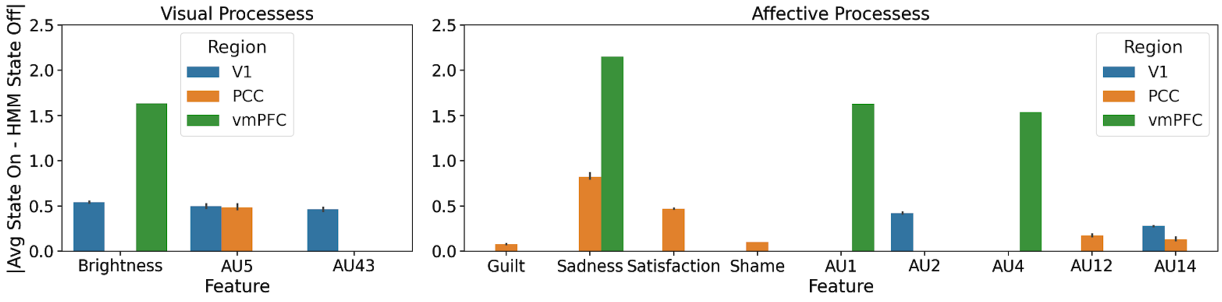

**Figure S16. Inter-Experiment Shared Response Model Face Expression Weights.** Here we visualize the transformation matrices learned from fitting the cross-experiment shared response model that project facial expressions into the latent components. The left column is a neutral face. The middle column depicts the vector field of how each facial landmark moves from a neutral position. The right column depicts the intensity of each facial action unit. Each row corresponds to a separate latent component.

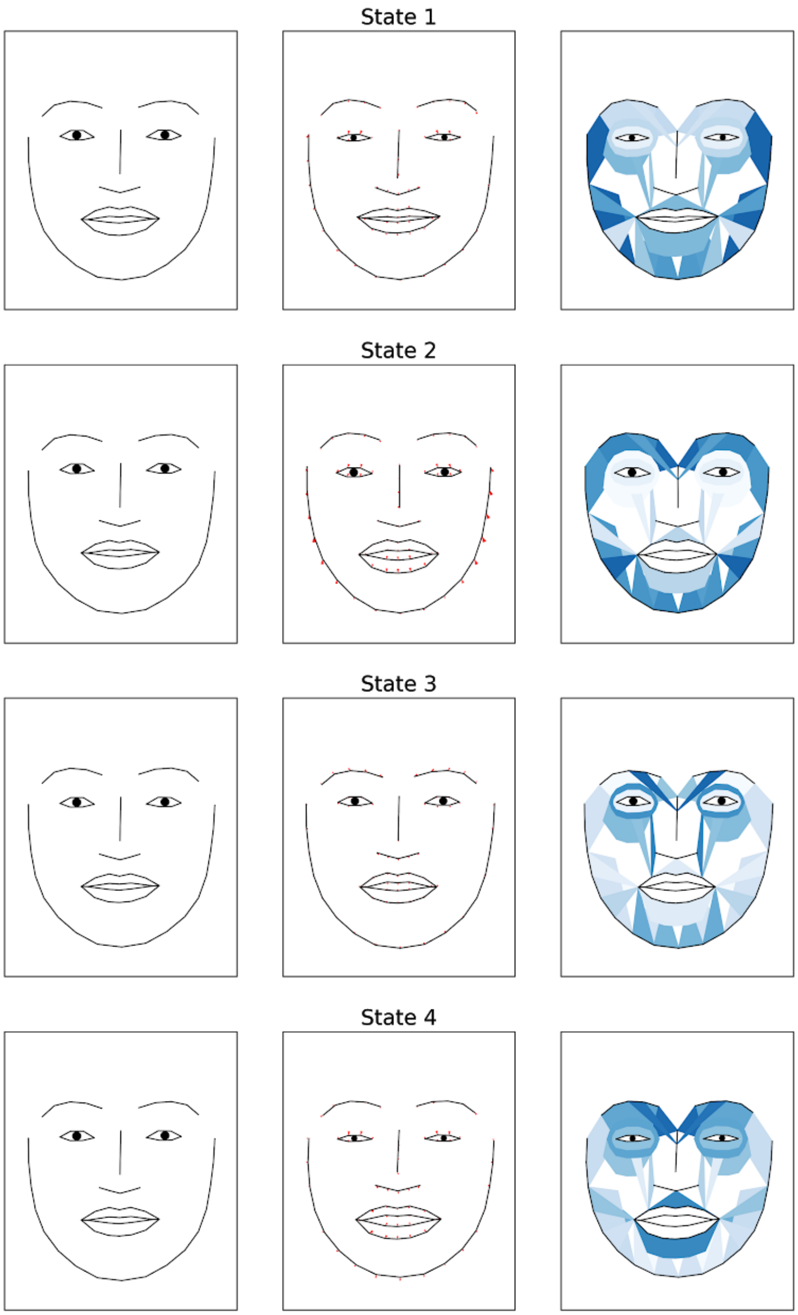

**Figure S17. Inter-Experiment Shared Response Model Emotion Rating Weights.** Here we visualize the transformation matrices learned from fitting the cross-experiment shared response model that project the self-reported subjective feelings into the latent components. Each panel corresponds to a separate latent component.

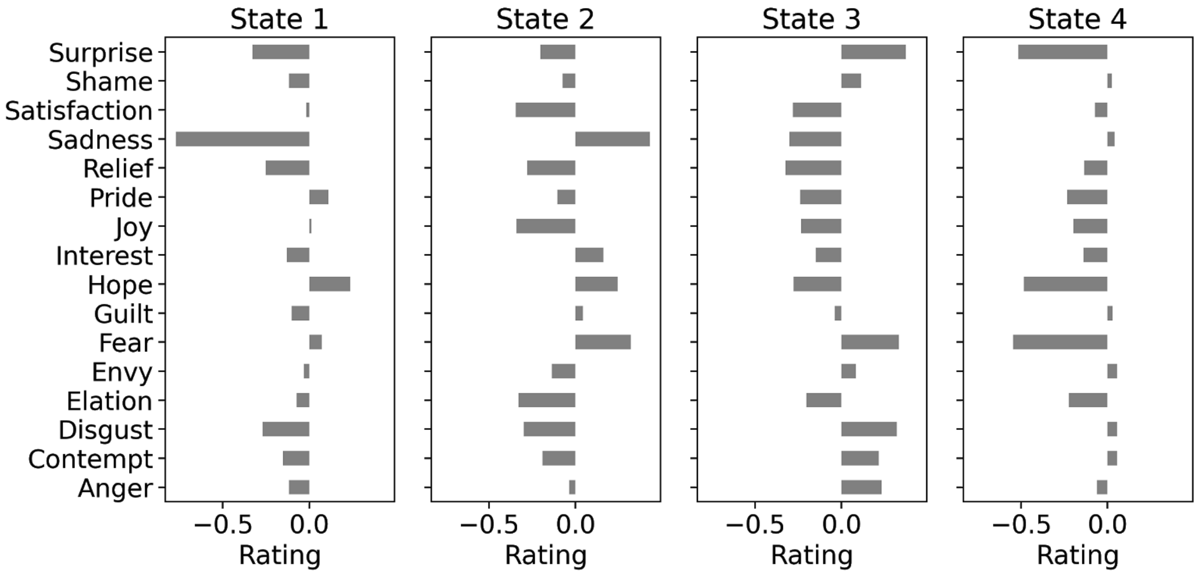

**Figure S18. HMM State Contrast Decoding.** Panel A depicts axial slice montages of whole-brain univariate contrasts that identify average activation changes when the vmPFC is in a particular state as indicated by the Group-HMM. Contrasts are thresholded using FDR q < 0.05. Panel B illustrates the top 10 Neurosynth reverse inference topic maps that correlate with each un-thresholded HMM State Contrast T-Map. Dotted circle indicates correlations r=0. Overall, we find that State 1 was most associated with maps predictive of Reward, Fear, Pain, Motor, and Somatosensory processing, State 2 was most associated with Social processing, and State 3 was most associated with Executive Control, Attention, Conflict, and Learning. The contrast maps can also be viewed on Neurovault (https://neurovault.org/collections/9062/).

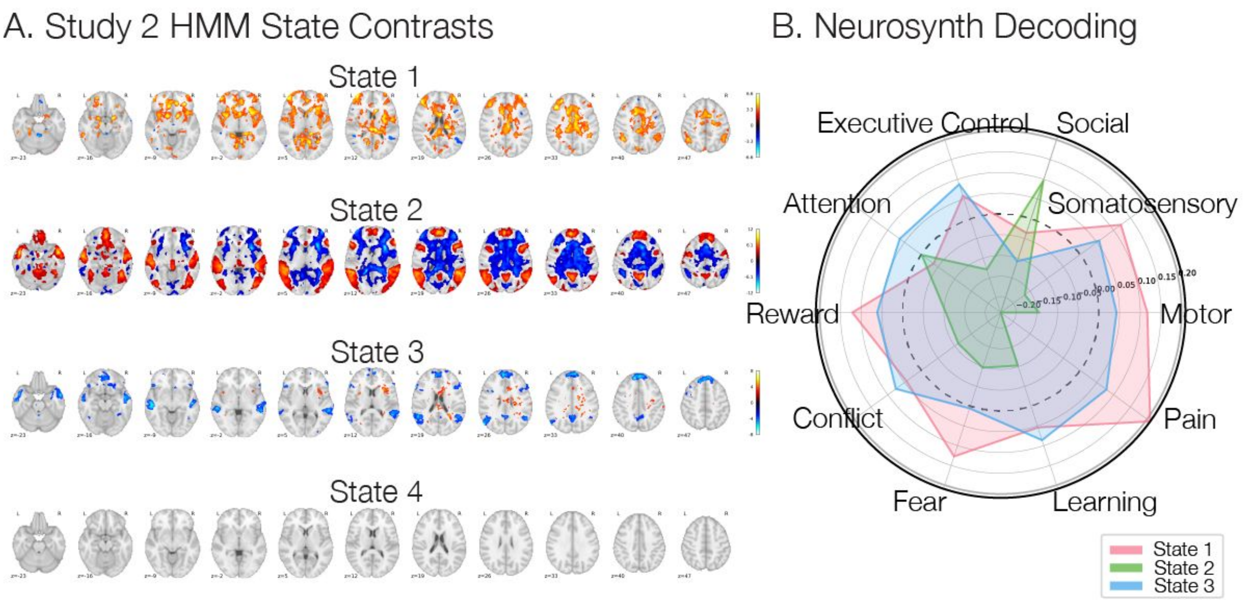
